# Mycobacterial HelD is a nucleic acids-clearing factor for RNA polymerase

**DOI:** 10.1101/2020.07.20.211821

**Authors:** Tomáš Kouba, Tomáš Koval’, Petra Sudzinová, Jiří Pospíšil, Barbora Brezovská, Jarmila Hnilicová, Hana Šanderová, Martina Janoušková, Michaela Šiková, Petr Halada, Michal Sýkora, Ivan Barvík, Jiří Nováček, Mária Trundová, Jarmila Dušková, Tereza Skálová, URee Chon, Katsuhiko S. Murakami, Jan Dohnálek, Libor Krásný

## Abstract

RNA synthesis is central to life, and RNA polymerase depends on accessory factors for recovery from stalled states and adaption to environmental changes. Here we investigated the mechanism by which a helicase-like factor HelD recycles RNA polymerase. We report a cryo-EM structure of an unprecedented complex between the *Mycobacterium smegmatis* RNA polymerase and HelD. The crescent-shaped HelD simultaneously penetrates deep into two RNA polymerase channels that are responsible for DNA binding and substrate delivery to the active site, thereby locking RNA polymerase in an inactive state. We show that HelD prevents non-specific interactions between RNA polymerase and DNA and dissociates transcription elongation complexes, but does not inhibit RNA polymerase binding to the initiation σ factor. The liberated RNA polymerase can either stay dormant, sequestered by HelD, or upon HelD release, restart transcription. Our results provide insights into the architecture and regulation of the highly medically-relevant mycobacterial transcription machinery and define HelD as a clearing factor that removes undesirable nucleic acids from RNA polymerase.

---

A smoothly functioning transcription machinery is essential for maintaining the physiologically relevant levels of gene products and adequate changes in transcription are necessary for cell survival when the environment changes. In bacteria, transcription is executed by a single enzyme, DNA-dependent RNA polymerase [RNAP; composition of the core enzyme: α_2_ββ’ω^1^]. The RNAP core is capable of transcription elongation and termination but not initiation. To initiate, a σ factor is required to form a holoenzyme that recognizes specific DNA sequences, promoters^2^. Besides the core subunits that are conserved in all bacteria some species contain additional subunits/associated proteins, such as δ and ε that are present in Firmicutes^3,4^, HelD that is present in Gram-positive bacteria^5^, or CarD and RbpA^6^ that are found in Actinobacteria. The smooth functioning of the transcription machinery then depends on concerted activities of RNAP and numerous transcription factors.

One of these factors is HelD^5^, a protein similar to SF1 helicases^7^ that associates with the RNAP core in the model gram-positive bacterium *Bacillus subtilis* (*Bsu*) where it was shown to be involved in transcriptional recycling^8^. HelD binds and hydrolyzes ATP and this is accompanied by conformational changes in the protein as demonstrated by SAXS experiments^9^. The absence of HelD from *Bsu* cells results in prolonged lag phase during outgrowth of stationary phase cells when diluted into fresh medium; overexpression of HelD then accelerates spore formation^10^. However, the structure of HelD, its binding mode to RNAP, and mechanistic details of its function are unknown.

Here, we present structural data for HelD from *Mycobacterium smegmatis* (*Msm*) in complex with the RNAP core and provide insights into its function. We solved the 3D structure of three complexes of *Msm* RNAP and HelD by cryogenic electron microscopy (cryo-EM). The structures represent a so far unknown type of interaction between an RNAP and a protein. Furthermore, we show that mycobacterial HelD and σ^A^ may simultaneously bind to the RNAP core and we provide biochemical evidence showing that in addition to ATP, HelD can also hydrolyze GTP. Finally, we demonstrate that HelD can both prevent binding of the RNAP core to non-specific DNA and actively remove RNAP from stalled elongation complexes.

## Results

### Cryo-EM of *Msm* RNAP-HelD complex

Our long-term attempts to crystalize *Bsu* HelD, RNAP core, or their complex failed; our cryo-EM experiments with the *Bsu* RNAP core were not successful; also, our recent SAXS-based data for the *Bsu* HelD-RNAP complex were not fully conclusive. However, in co-immunoprecipitation experiments with *Msm* RNAP we identified MSMEG_2174, a potential homolog of *Bsu* HelD (Figure S1). We also solved the X-ray crystal structure of *Bsu* HelD C-terminal domain (CTD), which was then used as a guide for building the model of *Msm* HelD.

We reconstituted a complex of *Msm* RNAP core and *Msm* HelD from purified recombinant proteins (Figure S2), and froze an isolated homogenous fraction of the complex on cryo-EM grids. We collected multiple preliminary cryo-EM datasets, which allowed us to optimize the cryo-EM conditions for high-resolution three-dimensional (3D) single-particle reconstructions (Figure S3). We identified two major 3D classes (State I and State II, Figure S4) at overall resolution ~3.1 Å (plus one subclass at ~3.6 Å), visualising almost the complete structure of HelD bound to the RNAP core in two conformations (Figure 1a,b, Movies 1,2), and one minor class (State III; Figure S4), at ~3.5 Å, which delineates only two domains of HelD binding to the RNAP core (Figure 1c, Movie 3). The structures of States I and II share the same overall fold of HelD, with a crescent-like shape. The main body of the crescent is sitting in between the domain 2 of the RNAP β subunit (also called β-lobe domain) and the funnel and cleft/jaw of the β’ subunit, burying about 774 and 2608 Å^211^ in State I and 1490 and 3623 Å^2^ in State II of binding surface area of β and β’ subunits, respectively. One end of the crescent protrudes deep into the primary channel, and the other end into the secondary channel of the RNAP core. Indeed, to be able to reach both RNAP channels simultaneously, the HelD protein is markedly elongated, around 200 Å along the outer edge of the virtual crescent, and the two ends of the HelD protein are separated by ~75 Å (State II; Figure 1b).

**Figure 1:**
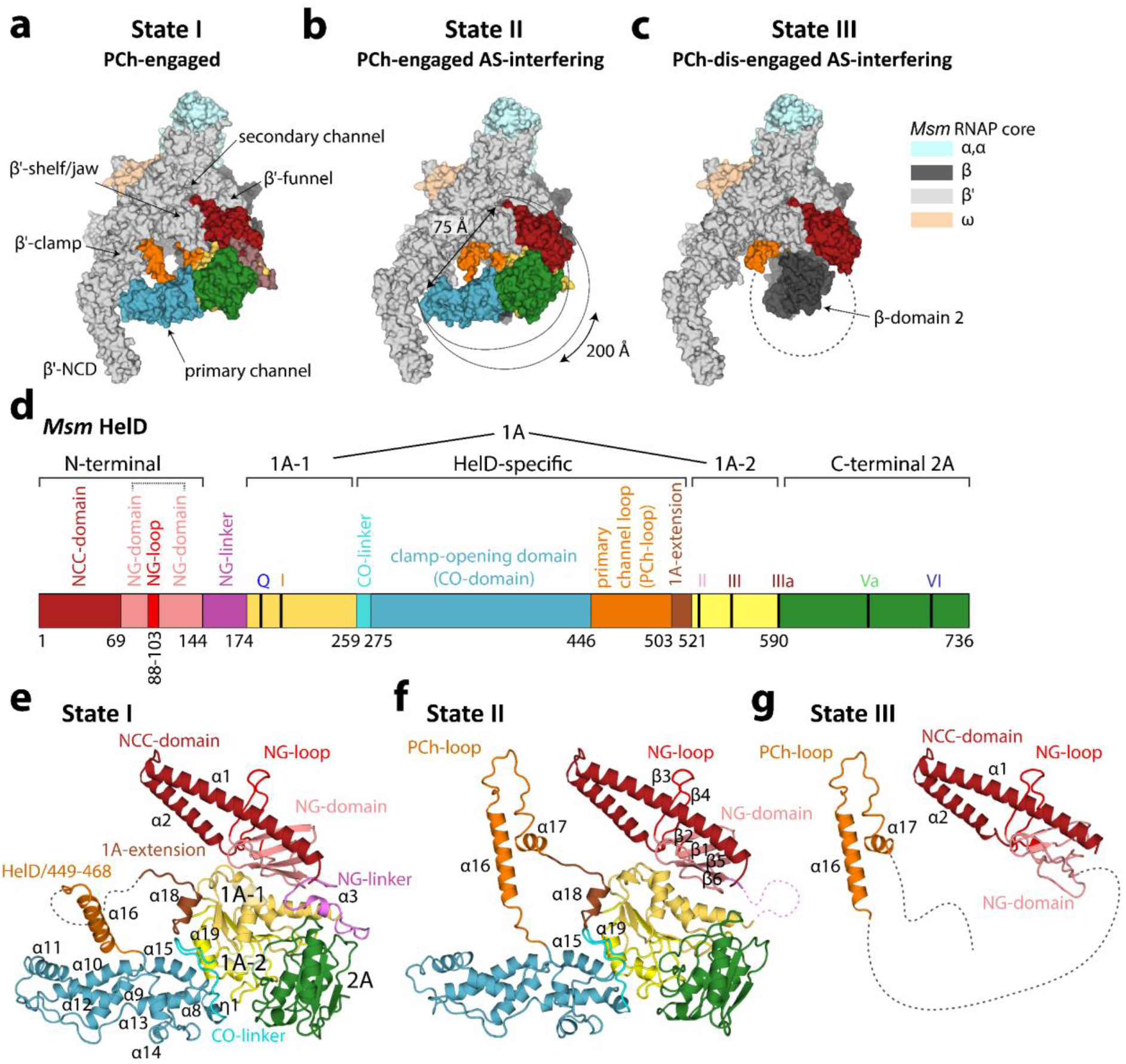
Cryo-EM structures of *Msm* HelD-RNAP complexes. **a, b and c**, Atomic model surface representation of three identified *Msm* HelD-RNAP complexes: State I – PCh-engaged, State II – PCh-engaged AS-interfering and State III – PCh-dis-engaged AS-interfering. When fully ordered in State I and II (**a** and **b**), the HelD protein (color coded as in **d**) forms a crescent-like shape, ends of which are protruding to the primary and secondary channel of the RNAP core. Partly ordered HelD protein in State III (**c**) vacates most of the RNAP primary channel. **d**, Schematic linear representation of the domain structure of the HelD protein. The 1A domain (two shades of yellow) is split in aa sequence into two parts, separated by a large HelD-specific insertion (hues of blue and orange). The nucleotide binding motifs are marked as vertical thick black lines. Aa numbering (*Msm*) is shown below. **e, f and g**, Three states of HelD as observed in **a, b and c** color coded according to the domain structure (**d**), secondary structure elements are marked as in Figure S5a.

The HelD protein itself is divided into six structured domains (Figure 1d-g), several of which possess unique novel folds. Interestingly, the 1A domain is composed of two parts (1A-1 and 1A-2) that are separated in the primary amino acid sequence by the intervening HelD-specific domain. According to the position of the HelD domains within the primary channel (PCh) and active site (AS), we name State I: PCh-engaged, State II: PCh-engaged and AS-interfering, and State III: PCh dis-engaged and AS-interfering (Figure 1a-c).

### The HelD N-terminal domain inserts into the RNAP secondary channel

The *Msm* N-terminal domain (HelD/1-144) forms an antiparallel α-helical coiled-coil (NCC) (HelD/1-69) followed by, and packed against the four-β-strand globular (NG) domain (HelD/70-144), which contains an additional prominent protruding loop (NG-loop, residues HelD/88-103, (Figure 1d-g and Figure 2a,b). The overall N-terminal domain structure is analogous to the archetypal fold interacting with the secondary channel of RNAP present in transcription factors such as GreA or ppGpp cofactor DksA^12–14^. Indeed, the HelD N-terminal domain interacts tightly with the secondary channel, burying ~1790 Å^2^ of interaction surface, contributing largely to the HelD-RNAP interaction. Several specific hydrogen bonds and salt bridges (Table S1a) are formed between the N-terminal domain and the secondary channel, and particularly the NG-loop specifically recognises the tip of the coiled-coil (CC) motif of the β’ funnel (Figure 2a).

**Figure 2:**
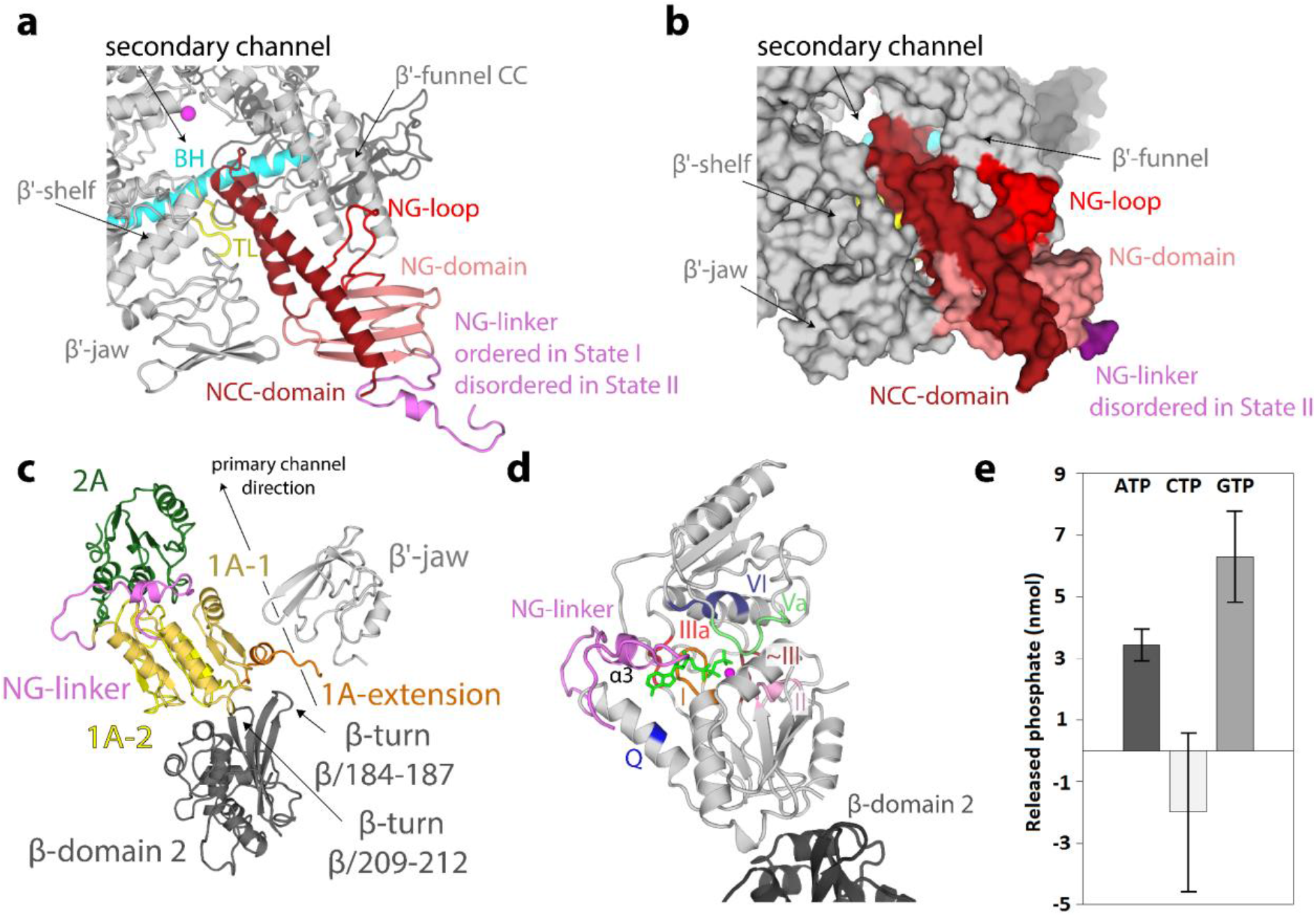
The HelD N-terminal domain inserts into the RNAP secondary channel; domains 1A-2A comprise the NTPase unit. **a** and **b,** Ribbon (State I) and surface (State II) representation of the HelD N-terminal domain interaction with the secondary channel of RNAP core (grey). The HelD coiled-coil domain (NCC-domain, firebrick) and the distinct loop (NG-loop, red) of the HelD globular domain (NG-domain, salmon) are inserted between β’ funnel, shelf and jaw. The NCC-domain reaches only the boundary line of the β’ bridge helix (BH, cyan) and leaves a passageway to the RNAP core active site (MgA, magenta sphere). The HelD NCC also restricts the trigger loop (TL, yellow) movement. The linker (NG-linker, violet) connects the N-terminal domain with domain 1A-1. **c**, The two *Msm* HelD Rossman fold domains (1A yellow and 2A green) form a canonical NTPase unit heterodimer with respect to structurally described SF1 helicases. Domain 1A tightly packs with β-domain 2 (dark grey) and its extension (brown) is clamped in-between one β-turn (β/184-187) of β-domain 2 and the tip of the β’ subunit jaw (light grey). **d**, Model of ATP binding to the conserved nucleotide binding site of motifs Q (blue), I (brown), II (pink), ~III (orange), IIIa (red), Va (palegreen) and VI (deepblue). ATP (green) and Mg^2+^ (magenta sphere) are added based on superposition with the ternary complex of UvrD (PDB ID 2IS4). **e**, HelD exhibits ATPase and GTPase activities but does not hydrolyse CTP.

The topology of the *Msm* HelD NCC is conserved in comparison with other secondary channel-interacting transcription factors (Figure S6); however, in contrast to the known structures of such complexes, the *Msm* HelD NCC is shorter and its tip does not reach into the AS (Figure S6). Indeed, a large part of the NCC is extensively packed with the NG-domain into a common hydrophobic core, thereby preventing the NCC to bind further towards the AS. The HelD NCC tip is positioned at the level of the RNAP AS β’ bridge helix (BH), ~10-12 Å away from Mg^2+^ metal A (MgA) of the AS, and as a result, it constitutes one wall of the secondary channel pore leading to the AS. The pore itself is approximately ~11 Å wide (Figure 2b) and this would still allow nucleoside triphosphate (NTP) passage into the AS. On the other hand, the NCC-domain restricts the conformational freedom and induces folding of the AS trigger loop (TL, β’/1009-1028). This would likely interfere with the nucleotide addition cycle.

Another difference with respect to GreA family transcription factors is that the HelD NCC tip does not contain the conserved DXX(E/D)^15–17^ motif (Figure S6), and it is therefore unlikely that the *Msm* HelD N-terminal domain possesses a Gre factor-like endonuclease activity.

### The NTPase unit of HelD is positioned in the vicinity of the downstream section of the primary channel

The presented structure confirms our previous prediction^9^ where HelD, similarly to SF1 helicases, RapA and UvrD, contains a conserved Rossmann fold 1A-2A heterodimer. 1A-1 is connected with the N-terminal domain by the NG linker (HelD/145-173), which orders only in State I. 1A-2 is then followed by 2A (Figure 1d,e,f and 2c).

The 1A domain docks on the β-domain 2 where it induces small domain-orientation and conformational changes and it prolongs the wall of the downstream section of the primary channel along the axis of the virtual downstream DNA (Figure 2c). The 1A domain buries an area of 725 Å^2^ of the interaction surface of the β-domain 2, the binding also involves ordering of the β-turn β/209-212 and many hydrogen bonds and salt bridges (Table S1b). In addition, the extension of the 1A domain (HelD/504-521) is clamped in between the prominent β-turn β/184-187 of the β-domain 2 and the tip of the β’ subunit jaw, further securing the 1A domain in its place.

The 1A-2A heterodimer establishes the canonical tertiary structure to form an NTP-binding pocket. Conserved residues of motifs Q, I, II, ~III, IIIa, Va, and VI are then likely involved in ATP binding^7,18^ (Figure 2d) while motifs and residues typical for DNA binding are missing. However, the base type specificity is not obvious from the structural data and therefore we measured nucleoside triphosphate hydrolysis activity of the isolated HelD protein. HelD showed strong hydrolysis activity of purine base nucleoside triphosphates but no activity towards a pyrimidine-containing counterpart (Figure 2e, S7f). We also added ATP or non-hydrolysable ATP analogue to the HelD-RNAP complex, but we were not able to visualize any NTP-bound state by cryo-EM. Indeed, the orientations of conserved HelD/Tyr589 and Arg/590 of motif IIIa, which are supposed to stack and coordinate the base and phosphate groups in the canonical ATP-bound state^19^, are incompatible with ATP binding in the HelD NTP-free state, both in State I and II (Figure S7a). Notably, helix α3 of the ordered NG-linker in State I covers the putative NTP-binding pocket and partially obstructs the site entrance (Figure S7a). However, the entire linker can become disordered as seen in State II (Figure S8h), which is probably more compatible with NTP binding (see details below).

The superposition of HelD 1A-2A with similar structures of UvrD (PDB ID 2IS4) (Figure S7b,c), PcrA (PDB ID 3PJR), AdnA/B (PDB ID 6PPR) and RapA (PDB ID 6BOG) confirms that the Rossmann fold domains are packed in the canonical mutual orientation. However, unlike in *bona fide* SF1 helicases^7,18^ where ssDNA is bound in the interface cleft of the dimer by conserved motifs Ia, Ic, IV, and V (Figure S7b,c), these motifs are not conserved in HelD. Instead, HelD contains proline-rich loops in place of these motifs and a large negatively charged surface patch in the equivalent areas (Figure S7d,e). Similarly, the ssDNA binding motifs are not conserved inRapA, a functional homolog of HelD, a helicase-like protein involved in recycling of RNAP. RapA, however, binds differently to RNAP than HelD^20^.

### The *Msm* HelD-specific domain is inserted into the downstream section of the RNAP primary channel

The HelD-specific insertion domain is composed of the clamp-opening domain (CO-domain, HelD/261-447) and the primary channel loop (PCh-loop, HelD/448-503) (Figure 1d, 3a,b). The CO-domain is an extended, mostly α-helical, and completely novel fold with no structure homologs (Figure S5b). On one side, the CO-domain packs against the 1A domain helix α19 and β-turn HelD/561-564. Additionally, the CO-1A interaction is stabilised by the CO-linker (HelD/259-275), which connects the two domains. In State I, the other side of the CO-domain, the CO-tip, butts against the three-stranded sheet of the β’ non-conserved domain (NCD) and an α-helix (β’/122-133) of the β’ clamp just preceding it (Figure 3a). The only significant ordered part of the PCh-loop in State I, the protruding helix α16 (HelD/451-468), is erected against the β’ three-stranded sheet (β’/1164-1210) and the α16 tip locks behind the helix-turn-helix motif β’/271-304 by HelD/Tyr466. Altogether, the α16 interaction with the β’ clamp might be helping the CO-domain insertion into the primary channel. In State II, the CO-domain fold alters and the PCh-loop completely refolds. The CO-tip shifts towards the β’ clamp coiled-coil domain (β’-CC) domain and reaches the peptide β’/387-389 of the rudder (Figure 3b). The PCh-loop protruding helix α16 refolds (α16 register slightly shifts towards the C-terminus of HelD) and dis-engages with the β’ three-stranded sheet (β’/1164-1210), and the whole PCh-loop orders towards the AS (see next section). Correspondingly, the two insertion modes of the CO-domain and PCh-loop into the primary channel force the β’ clamp domain to swing out into two distinct positions (see details below).

**Figure 3:**
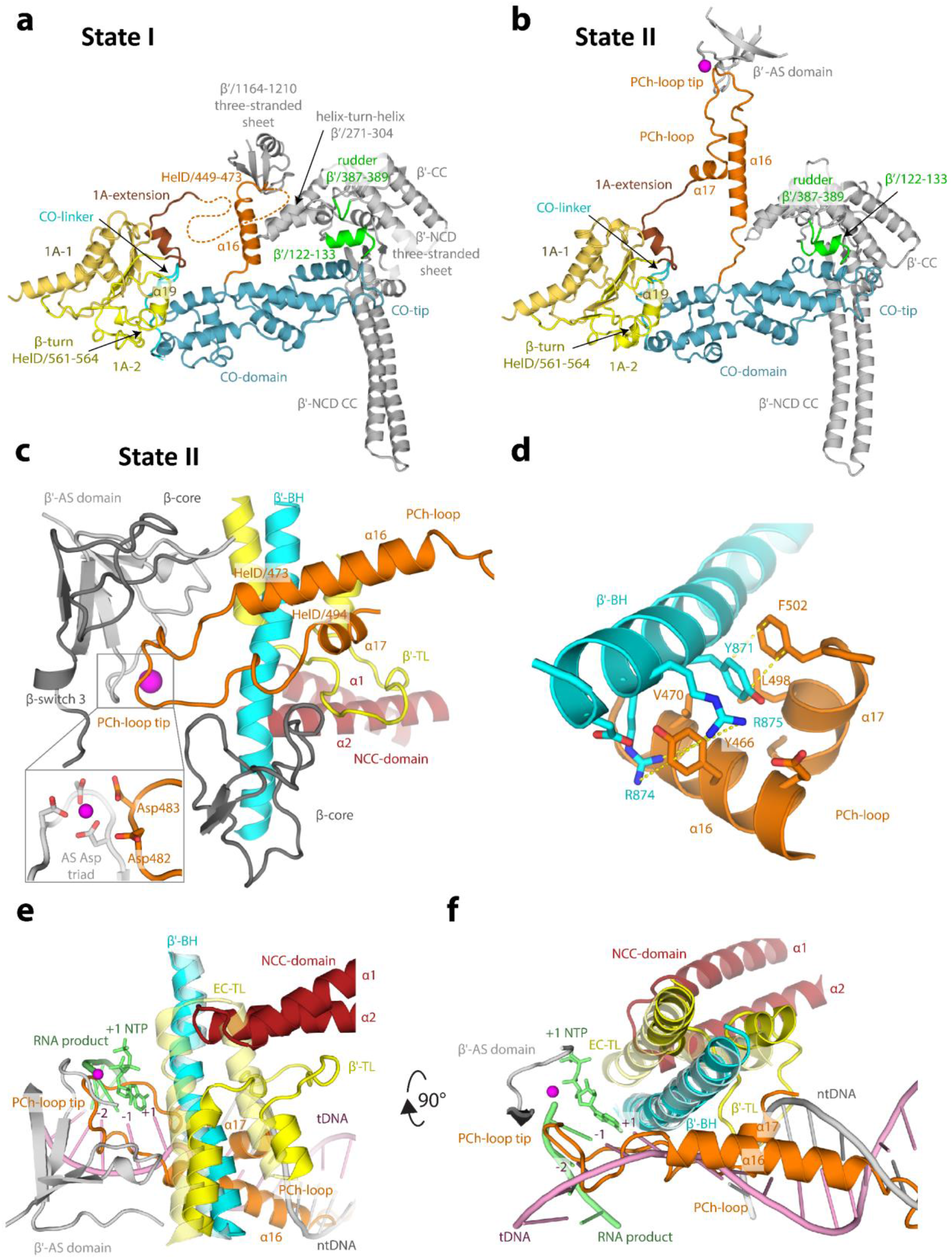
The *Msm* HelD-specific domain interactions with the RNAP primary channel. **a** and **b,** Ribbon representation of the HelD-specific domain inserting into the RNAP primary channel in State I (**a**) and State II (**b**). In State I (**a**), the clamp-opening (CO, blue) HelD-specific domain is projected from the HelD 1A domain (yellow) towards the β’ clamp (grey). At one end, the CO is bonded to the 1A domain by the CO-linker (cyan), and stabilised by β-turn 561-563 and α19 (yellow). On the other end, the CO-domain tip abuts towards the β’ NCD three-stranded sheet. Concomitantly, the HelD helix α16 (part of peptide HelD/449-473, orange) butts against the β’/1164-1210 three-stranded sheet. The connection between α16 and the 1A-extension is disordered (dotted line). In State II (**b**), The CO interaction with the 1A domain remains similar to State I (**a**). The CO-domain tip, however, shifts towards the β’ rudder (green) and β’/122-133 α-helix. Concomitantly, the HelD PCh-loop (orange) folds towards the active site (MgA, magenta sphere) and folds back towards the 1A-extension (brick) and 1A domain. **c**, The PCh-loop folds into the RNAP active site. The HelD loop 473-494 and the two adjacent α-helices (α16 and α17, orange) fold alongside the RNAP bridge helix (BH, cyan) towards the RNAP active site and HelD/Asp482 directly contacts the MgA (magenta sphere, details in inset). The RNAP trigger loop (TL, yellow) is restricted and folded between the HelD PCh-loop helix α17, the HelD NCC-domain (ruby), β’ BH and the β core domain (dark grey). **d**, Detail of the BH interaction with HelD α16 and α17. BH β’/Arg874 and 875 sandwich HelD/Tyr466, and β’/Tyr871 stacks on HelD/Phe502. The hydrophobic interactions are marked by yellow dotted line. **e** and **f**, The HelD PCh-loop binding in the active site chamber is mutually exclusive with the presence of the transcription bubble. Two perpendicular views of superposition of the *Tt* RNAP elongation complex (PDB ID 2O5J, pale colors) and HelD State II (solid colors) are shown. The folded TL in pre-translocated EC would sterically clash with the HelD NCC-domain. The HelD PCh-loop tip would sterically clash with RNA/DNA hybrid at positions +1 to −2, and the HelD α16 and α17 helices would clash with downstream DNA duplex. Color code as in **3c**, template DNA in pink, non-template DNA in grey, product RNA and incoming NTP at position +1 in green.

### The HelD PCh-loop is able to fold into the RNAP active site

In the cryo-EM map of the AS-interfering State II, high-resolution density is present for the entire register of the PCh-loop, which is folded in the AS cavity of RNAP (Figure 3c,d and S3f). The folding of the PCh-loop in-between the walls of the AS chamber is also compatible with the regular open form of the RNAP core as observed in State III.

In comparison to State I, in State II the protruding helix α16 refolds, the helix register shifts to residues 455-472, and together with a newly folded helix α17 (HelD/495-500) they tightly pack with the second half of the β’ BH (Figure 3d). In detail, BH β’/Arg874 and 875 sandwich α16 HelD/Tyr466 and, cooperatively, BH β’/Tyr871 stacks on HelD/Phe502 and is inserted into a hydrophobic pocket formed by HelD/Tyr466, Ala467, Val470, and Leu498. The rest of the PCh-loop (HelD/473-494) specifically wedges into the AS cavity (Table S1c), towards the AS aspartate triad and the MgA. Notably, there are four acidic residues (482-DDED-485) at the very tip of the PCh-loop and the HelD/481-483 peptide folds along the AS β-strand β’/537-544, such that HelD/Asp483 is in contact with MgA and HelD/Asp482 in its near proximity (Figure 3c and S3f). HelD/Asp482 interacts with β’/Arg500, HelD/Glu484 stabilizes the loop in the active site by interaction with β/His1026, and HelD/Asp485 contributes to the AS-interfering loop stability by a salt bridge with the side chain of HelD/Arg477. Two other motifs support formation of the PCh-loop structure in the RNAP AS – a small hydrophobic core formed by the HelD/Val475, Leu480, and Leu488 side chains and an intra-chain ion-pair HelD/Arg477-Asp491, with HelD/Arg477 leaning against β/Pro483.

As a result of the PCh-loop folding into the primary channel and HelD NCC folding in the secondary channel, the NCC tip and the tip of the PCh-loop are brought close together (the distance is only 17 Å). This also restricts the trigger loop, which is therefore partially folded in the space between BH, HelD α_2_ and α17, the peptide between α17 and α18, and the peptide of β/Ile182-Glu187. In summary, the PCh-loop seems to interfere with the AS cavity so that it is not compatible with the NTP addition cycle. Moreover, the superposition with a model of *Thermus thermophilus* (*Tt*) RNAP EC (PDB ID 2O5J) (Figure 3e,f) suggests that the whole PCh-loop would be in steric clash with the dwDNA duplex and the RNA/DNA hybrid in the AS as far as position −2. A parallel can be drawn between the presence of the PCh-loop in State II and the so called DNA-mimicking loop of PolI^21^, which also occupies the AS chamber and the surroundings of the AS and is sterically incompatible with the existence of the DNA transcription bubble in RNAP.

### Global domain changes of RNAP upon HelD binding

Superposition based on the β-core region (β/430-738) of the *Msm* RNAP core (PDB ID 6F6W), elongation complex (EC, based on PDB ID 2O5J) and States I-III enables analyses of global differences of the three observed structural states (Figure S8). In all States, when compared to the model of EC structure, probably just the insertion of the HelD N-terminal domain into the secondary channel slightly alters the position of the β’-jaw/shelf and β-domain 2 (Figure S8g) and this alteration may weaken interaction with dwDNA, reminiscent of TraR (a distant DksA homolog) binding to *E. coli* RNAP^22^

The major change between the States is the interplay between the refolding of the PCh-loop and the CO-domain position in the primary channel. In State III, solely the PCh– loop’s tight contact with the AS stabilizes a very open form of RNAP (Figure S8a,b,f), ~33 Å at the narrowest point of the primary channel (measured by the distance of the Cα atoms of β/Lys273 and β’/Lys123), comparable to the structures Core2 and Holo2^1^ (32.2 and 33.6 Å, respectively). In State I, the PCh–loop’s interaction with β’ helix-turn-helix and three-stranded sheet, and the CO-domain insertion into the primary channel make the opening of the RNAP clamp (~35 Å; Figure S8a,b) slightly wider than the extremely open forms of the Lipiarmycin-(PDB ID 6FBV^23^) and Fidaxomicin-locked (PDB ID 6C06^24^) RNAPs (34.2 and 33.6 Å, respectively^1^). In State II (Figure S8e), while the CO-domain still inserted, the PCh-loop abolishes the β’ contact and folds in the AS instead, and this forces the β’-clamp (β’/1-406) to rotate with respect to the remaining parts of the complex so that the β’-NCD CC tip opens further away from the juxtaposed β-domain 2 but at the same time the β’-rudder, β’-CC and adjacent secondary elements move about 11 Å closer to the tip of the HelD CO-domain. The RNAP clamp is therefore splayed by unprecedented 45 Å (Figure S8b) and this likely facilitates DNA release.

The next major difference are the β-domain 2 and CO-domain adjustments upon change of the 1A-2A heterodimer (Figure S8e). The mutual orientation of 1A and 2A domains between States I and II is almost preserved, although with much poorer density for 2A in State II. This most likely stems from the more pronounced mobility of 2A, possibly linked with the lack of stabilization by the unfolded NG-linker in State II. The 2A relaxation allows movement of 1A in respect to the N-terminal domain (~3° difference measured by HelD α1 and α5) and a concomitant shift of both the β-domain 2 and CO-domain (Figure S8e). In detail, this global change is accompanied by a shift and changes in the secondary structure of HelD/230-252 within the 1A domain (largest shift about 9.3 Å for Val245). Helix α6 is extended and helix α7 is formed in State II (Figure S5a) and 1A-extension shifted. State I interactions between α6 and the NTPase site, and α6 and the NG-linker that are NTP-binding prohibitive, are broken in State II and the NTPase site of HelD becomes wide open (NTP-binding permissive; Figure S8h). Although this change makes the NTPase site accessible for NTPs, additional conformational changes are still required for NTP accommodation.

Finally, HelD binding in States I and II also leads to opening of the RNA exit channel between the β-flap and β’-lid and β’-Zn-finger by about 15 Å and 21 Å, respectively (Figure S8c,d). State III keeps the channel still rather open by about 12 Å. This is expected to contribute to RNA release.

### HelD clears the RNAP primary channel

The position of the HelD CO-domain in the primary channel of RNAP suggests that HelD may prevent non-specific interactions between the RNAP core and DNA. To test this, we performed electrophoretic mobility shift assay (EMSA) with RNAP and a fragment of mycobacterial DNA in the presence/absence of HelD. Figure 4a,b,c shows that HelD significantly abolishes the nonspecific binding of the RNAP core to DNA.

**Figure 4:**
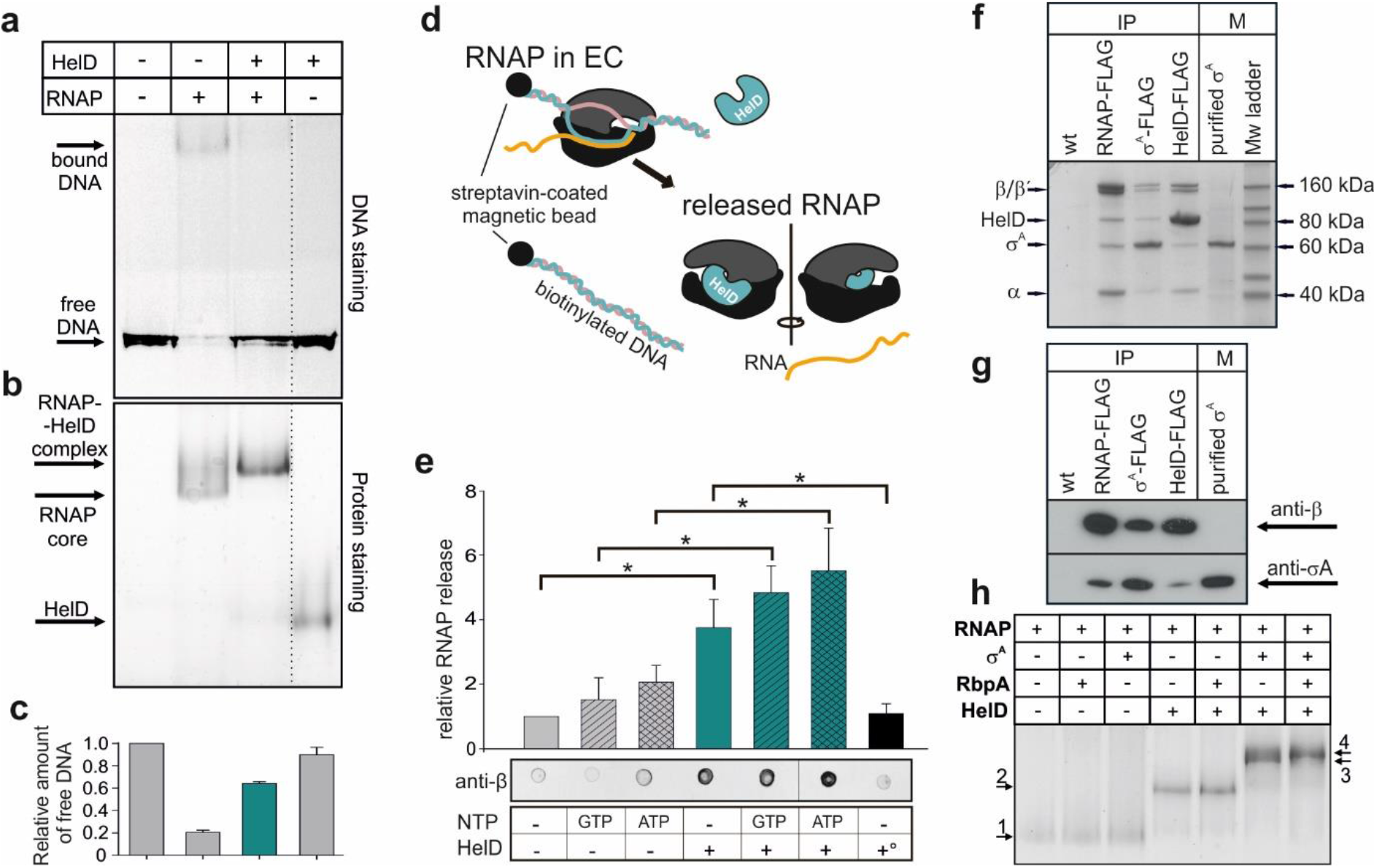
Binding of *Msm* HelD to RNAP and its effects on DNA-RNAP interactions. **a**, DNA binding to RNAP - EMSA - binding of 300 bp DNA to the *Msm* RNAP core and the effect of HelD. **b,** the same gel as above but stained for proteins. The dotted line shows where the gel was electronically assembled. **c**, quantitation of EMSA– the bars here (the amount of unshifted DNA) and in **e** are averages from at least three independent experiments, the error bars show ±SD. **d**, EC disassembly - scheme: ECs were assembled on DNA:RNA scaffolds and challenged with HelD and/or NTPs. RNAP released into buffer was quantitated by Western dot blots. **e**, Quantitation of EC disassembly experiments. Representative primary data are shown below the graph. Presence/absence of individual components is indicated. +° indicates heat-inactivated HelD. The statistical significance in **e** for the indicated combinations was p<0.05 (asterisks). **f**, SDS PAGE of immunoprecipitations of *Msm* RNAP (β), σ^A^, and HelD. All proteins were FLAG fusions, the antibody was anti-FLAG. Wt, a strain without any FLAG fusion. The identity of the bands was confirmed by mass spectrometry. IP, immunoprecipitation; M, markers. **g**, Western blot of IPs of FLAG-tagged *Msm* RNAP (β), σ^A^, and HelD. Antibodies against RNAP β and σ^A^ were used to detect the presence of proteins in complexes. M, marker. **h**, *In vitro* protein interactions - EMSA. Proteins were detected by Simply blue SafeStain. In all cases, RNAP was first reconstituted with HelD and then with RbpA and/or σ^A^. A small, but reproducible shift was observed after addition of both RbpA and σ^A^ to RNAP-HelD, indicating the presence of all proteins in one complex. Numbered arrows indicate complexes with different protein composition (determined by mass spectrometry). In some cases, complexes with different protein composition displayed the same migration in the gel: 1. RNAP, RNAP-RbpA; RNAP-σ^A^; 2. RNAP-HelD, RNAP-HelD-RbpA; 3. RNAP-HelD-σ^A^; 4. RNAP-HelD-σ^A^-RbpA.

Moreover, we speculated that HelD might not only prevent DNA binding, but also actively disassemble stalled ECs. Stalled ECs (due to *e. g.* damaged DNA) are obstacles for both the coupled transcription-translation machinery^25,26^ and also for replication^27^, with potentially deleterious consequences if not removed. To test the ability of HelD to rescue stalled RNAP, we assembled ECs with the RNAP core on a DNA-RNA scaffold and challenged them with HelD in the presence/absence of NTPs (Figure 4d). HelD then, relative to mock treatment, was able to disassemble stalled ECs (Figure 4e). This process, interestingly, appeared to be independent of ATP or GTP.

### HelD, σ^A^ and RbpA can simultaneously bind RNAP core

Next, we tested the mode of binding of HelD to RNAP, asking whether the HelD-RNAP complex is compatible with the presence of other factors. Immunoprecipitation (IP) and Western blot experiments with FLAG-tagged *Msm* RNAP revealed the presence of HelD and σ^A^ (Figure 4f,g); FLAG-tagged *Msm* σ^A^ pulled down the RNAP core and HelD; FLAG-tagged HelD pulled down the RNAP core and σ^A^. These results suggested but not proved that HelD, σ^A^, and RNAP could be in one complex. Alternatively, HelD and σ^A^ could bind each other independently of RNAP. To decide between the two possibilities, we first pulled down FLAG-tagged HelD and associated proteins and from this mixture we subsequently pulled down FLAG-tagged σ^A^. Figure S9 shows the presence of HelD and RNAP in the second pull-down, demonstrating that these proteins can coexist in one complex. Additionally, RbpA, albeit in low amounts, was also present in the HelD-immunoprecipitated complex and RbpA-FLAG pulled down RNAP with σ^A^ and HelD (Figure S10). We then confirmed the interactions between the RNAP core, σ^A^ RbpA, and HelD by *in vitro* EMSA (Figure 4h). Modelling of hypothetical complexes of RNAP-HelD with σ^A^ and RbpA then revealed that relatively small changes in conformations of these proteins could allow their simultaneous binding to RNAP in States I-III (Figure S11 and Discussion).

## Discussion

This study describes a structurally unique complex between *Msm* RNAP and the HelD protein, defines its DNA-clearing activity, and outlines its role in transcription.

Previous biochemical studies used HelD from *Bsu*, which is only 21 % identical with the *Msm* homologue. Selected sequence homologs of *Msm* HelD are shown in Figure S12, revealing two main differences between *Msm* (Actinobacteria) and *Bsu* (Firmicutes). The first marked difference is the absence of ~30 aa from the N-terminal NCC-domain region in *Msm* HelD. This is consistent with the *Bsu* HelD NCC-domain protruding much deeper into the RNAP secondary channel and even overlapping with the AS (See accompanying papers Newing *et al.,* 2020; Pei *et al.,* 2020). The other difference is in the HelD-specific region where *Bsu* HelD completely lacks the PCh-loop. On the other hand, the organisation of the 1A-1 and 1A-2 split followed by the complete 2A domain is maintained (Figure 1d,e,f).

Based on the structural and functional data we propose a role for *Msm* HelD in transcription (Figure 5). We envisage that upon transcription termination when RNAP fails to dissociate from nucleic acids^28^, or in the event of stalled elongation, *Msm* HelD first interacts with RNAP by its N-terminal domain, likely competing for binding to the secondary channel with GreA-like factors. This initial HelD binding induces changes in β-domain 2 and β’-jaw/shelf (Figure S8g), possibly leading to destabilisation of DNA in the primary channel. The trigger loop is conformationally locked. Subsequently, the CO-domain and PCh-loop approach the primary channel. The PCh-loop, which is probably flexible in the RNAP-unbound state, folds partially upon binding RNAP (captured in State I) and then it penetrates deep into the primary channel, fully folds, and binds to the AS (captured in States II and III). The CO-domain interactions with β’ then secure the primary channel wide open (Figure S8a,b). At the same time, the RNA exit channel dilates (Figure S8c,d). All these processes lead to expulsion of any contents of the AS (compare states within Figure S8a).

**Figure 5:**
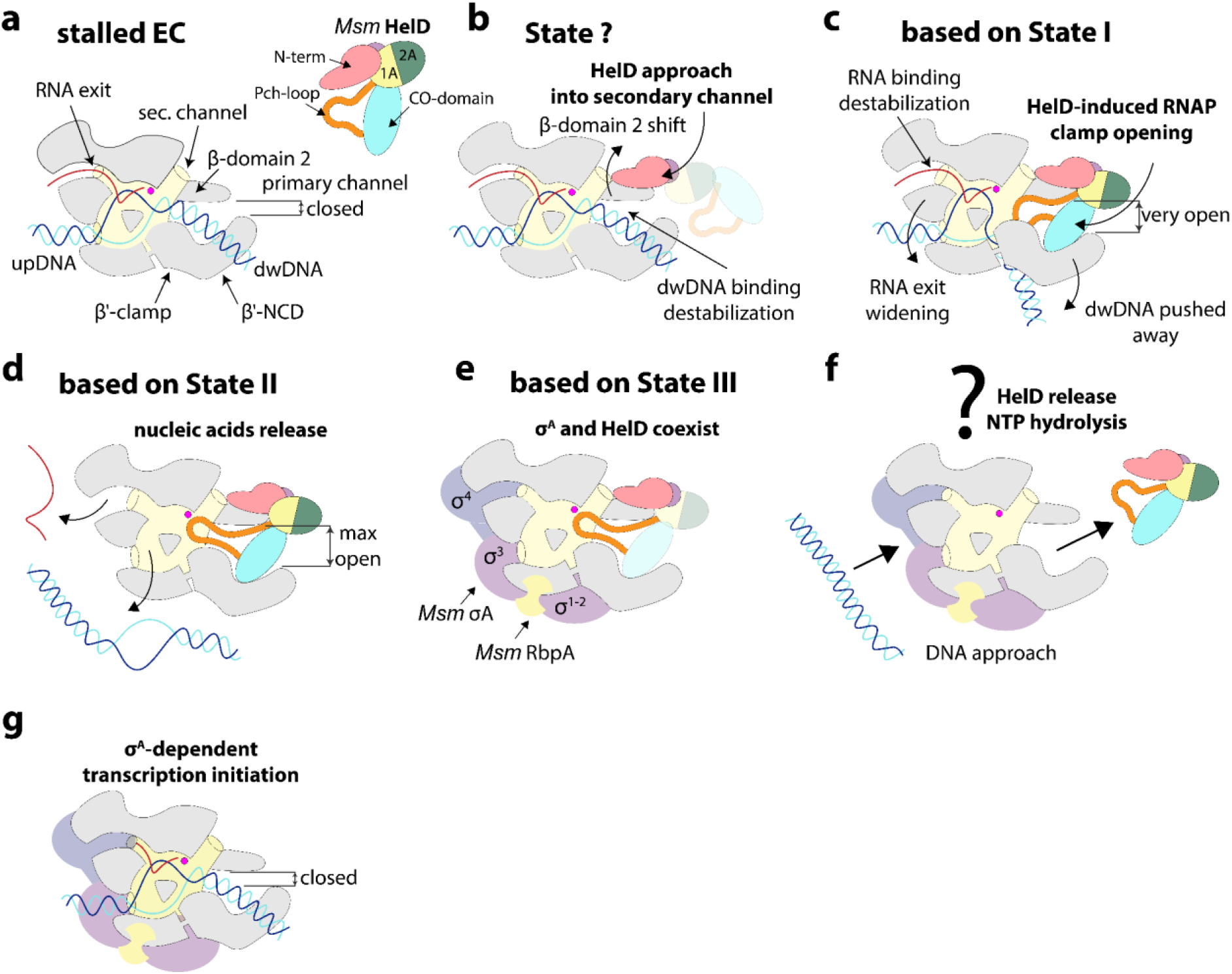
Schematic diagram of HelD function in RNAP recycling. **a**, When EC stalls and needs to be disassembled, **b** the HelD N-terminal domain (pink) first approaches the RNAP secondary channel and then induces changes in RNAP likely destabilising the dwDNA interaction. **c**, Subsequent interaction of HelD PCh-loop (orange) and the whole HelD-specific domain (cyan) in the RNAP primary channel open the RNAP cleft, widens the RNA exit channel and mechanically interferes with dwDNA. **d**, Even broader cleft/RNA exit opening together with the PCh-loop intervening deep in the AS (MgA, magenta sphere) displace dwDNA and the RNA/DNA hybrid from the active site cavity. **e**, The HelD-RNAP nucleic acid-free complex binds σ^A^ factor and RbpA, and all factors can bind RNAP core simultaneously. **f**, The complex binds to DNA promoter via σ^A^ factor with concomitant displacement of HelD from RNAP by an unknown mechanism, possibly dependent on NTP hydrolysis by HelD and **g**, a new round of σ^A^-dependent transcription cycle can initiate.

We note that neither HelD loading onto RNAP nor RNAP clamp opening nor EC disassembly are dependent on NTP hydrolysis. Energy from NTP hydrolysis is probably required to release HelD from its tight contact with RNAP. Free energy corresponding to ATP hydrolysis under physiological conditions in cells is around –50 kJ/mol ^29^. This is comparable to the estimated desolvation energy of the HelD:RNAP core interaction of –33.5 kJ/mol (Δ^i^G) for State I and –57 kJ/mol for State II. However, States I and II are not fully compatible with canonical NTP binding in the HelD NTPase unit. It remains to be answered which structural changes are required to actually enable NTP binding and hydrolysis.

Interestingly, HelD, σ^A^, and RbpA can co-occur on RNAP (Figures 4h and S9-11). This differs from *Bsu* where simultaneous HelD and σ^A^ binding has not been detected^8^. Regardless of the exact mutual σ^A^, RbpA and HelD positions, RNAP must subsequently assume a conformation that is compatible with promoter DNA binding and transcription initiation.

To summarize, HelD clears RNAP of undesirable nucleic acids, which likely contributes to the smooth functioning of the transcription machinery. Furthermore, it is conceivable that HelD may also function similarly to 6S RNA^30^ or Ms1^31^, keeping RNAP in an inactive state under unfavourable conditions. This stored RNAP then accelerates restart of gene expression when conditions improve.

Finally, the RNAP-inactivating ability of HelD might be utilized in development of specific antibacterial compounds that would stabilize the non-productive HelD-RNAP complex, shifting the equilibrium of RNAP states towards effective transcription inhibition, as seen *e.g.* in the action of fidaxomicin towards *M. tuberculosis* RNAP in complex with RbpA^24^.

## Supporting information

Msm_HelD_RNAP stateI

Msm_HelD_RNAP stateII

Msm_HelD_RNAP stateIII

## Methods

### Bacterial strains and plasmids

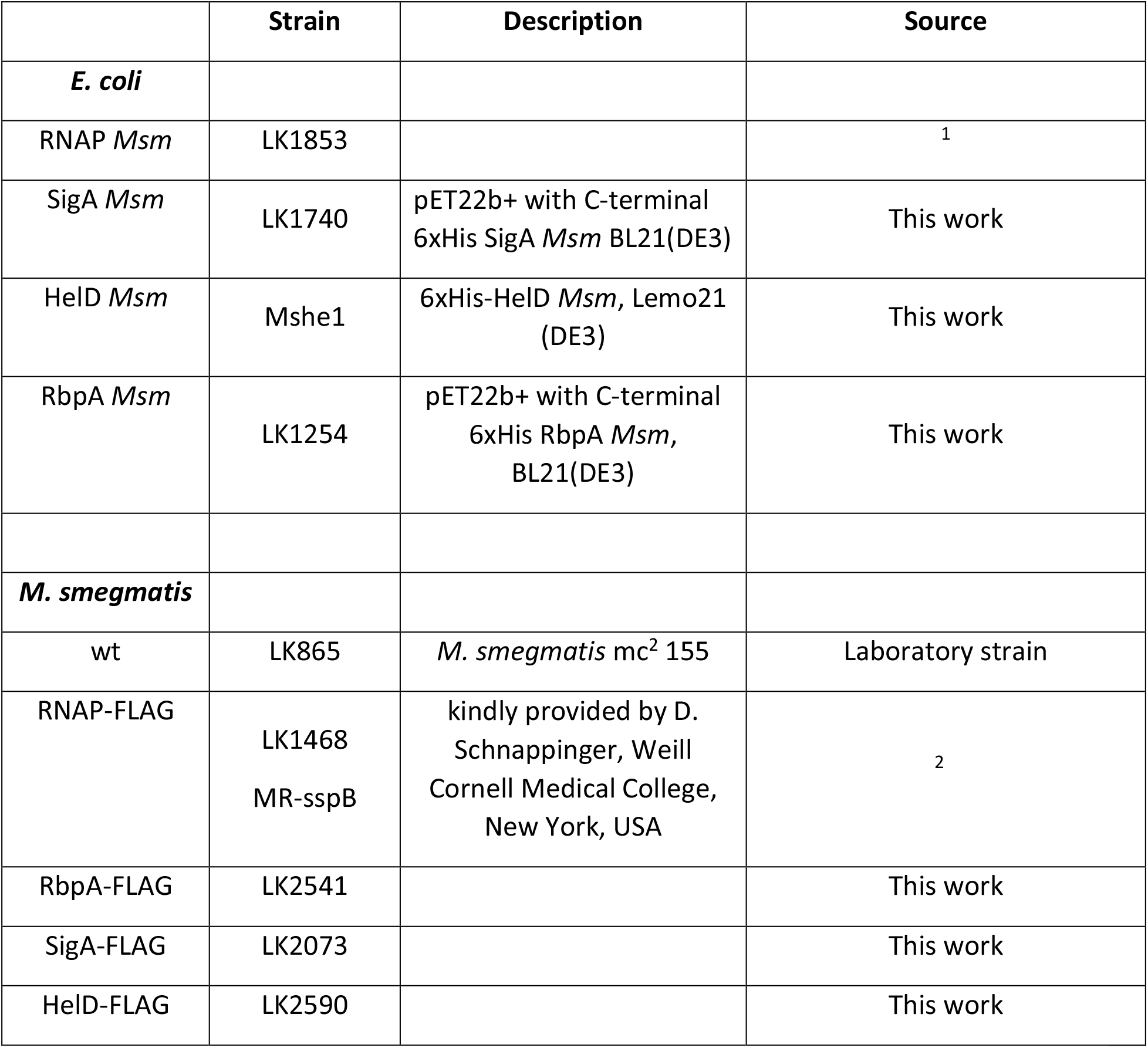

|  | Strain | Description | Source |
| --- | --- | --- | --- |
| ***E. coli*** |  |  |  |
| RNAP *Msm* | LK1853 |  | 1 |
| SigA *Msm* | LK1740 | pET22b+ with C-terminal 6xHis SigA *Msm* BL21(DE3) | This work |
| HeID *Msm* | Mshe1 | 6xHis-HeID *Msm*, Lemo21 (DE3) | This work |
| RbpA *Msm* | LK1254 | pET22b+ with C-terminal 6xHis RbpA *Msm*, BL21(DE3) | This work |
| ***M. smegmatis*** |  |  |  |
| wt | LK865 | *M. smegmatis* mc<sup>2</sup> 155 | Laboratory strain |
| RNAP-FLAG | LK1468<br>MR-sspB | kindly provided by D. Schnappinger, Weill Cornell Medical College, New York, USA | 2 |
| RbpA-FLAG | LK2541 |  | This work |
| SigA-FLAG | LK2073 |  | This work |
| HeID-FLAG | LK2590 |  | This work |

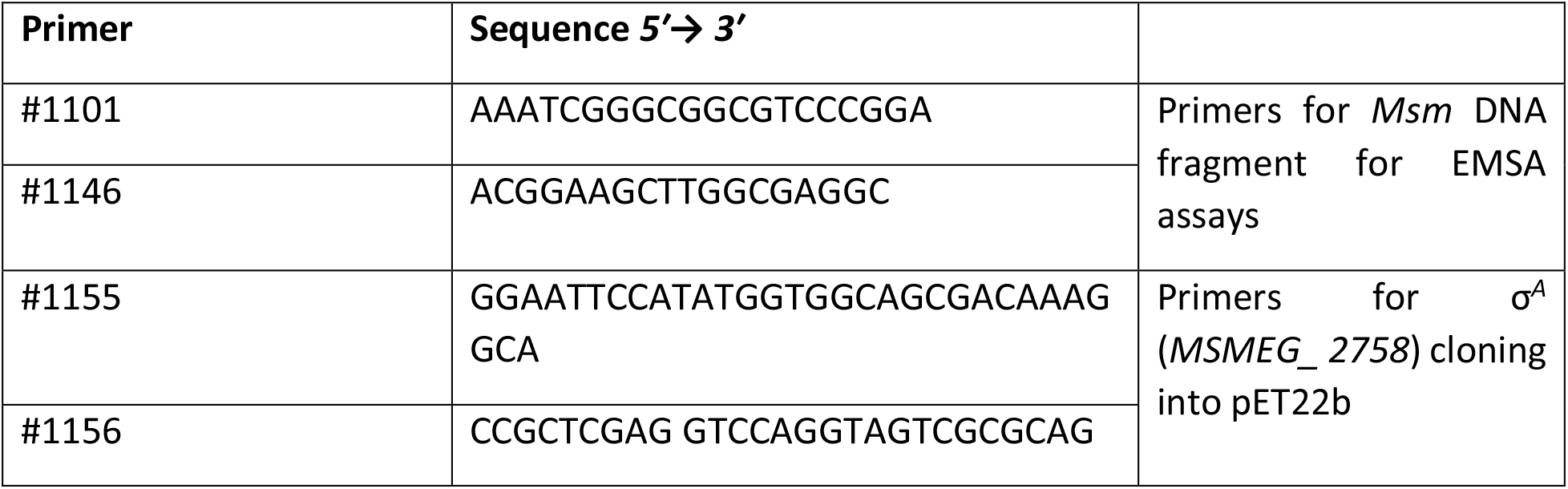
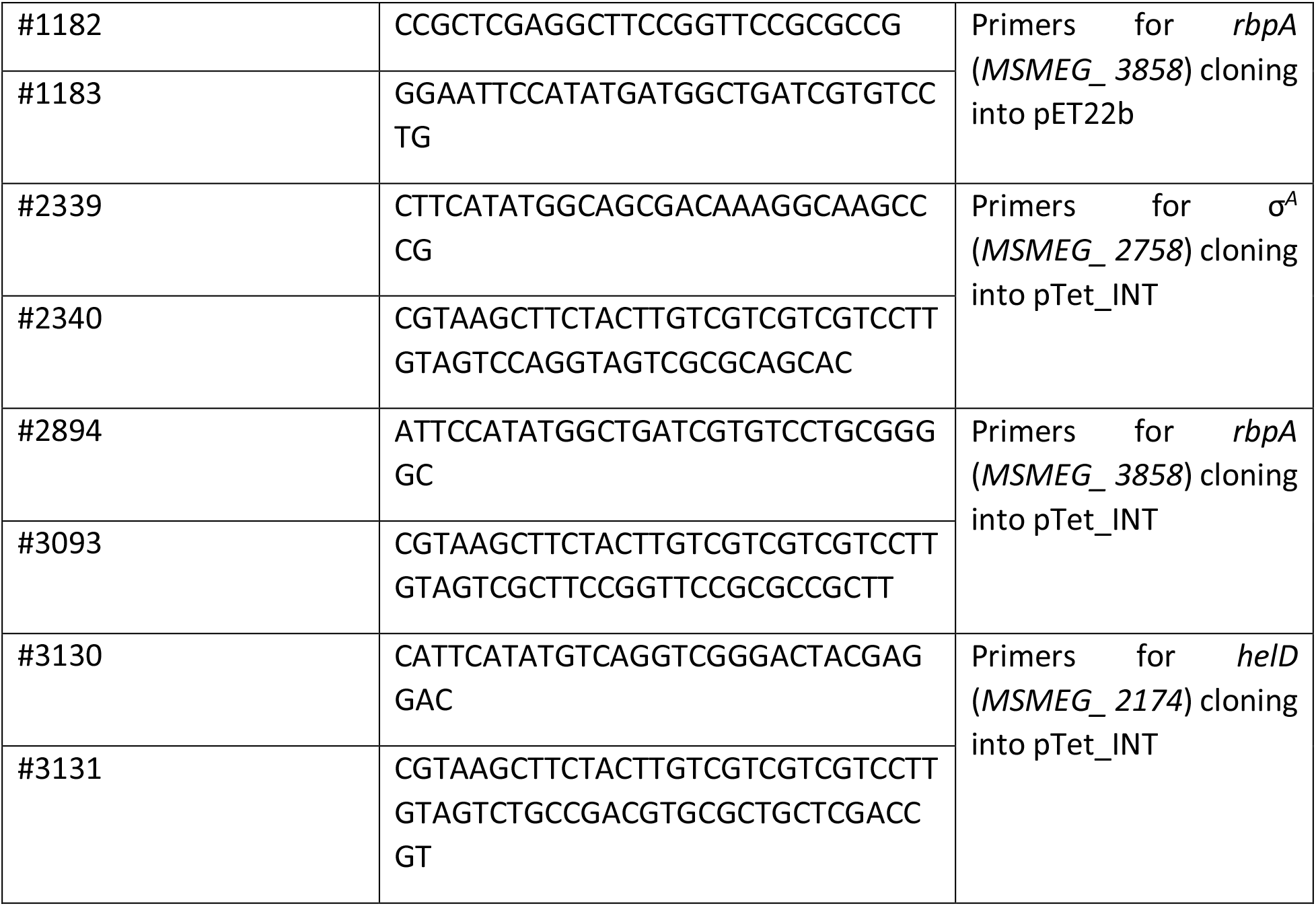

### Strain construction

#### σ^A^ and RbpA

σ^*A*^ (*MSMEG_2758*) and *rbpA* (*MSMEG_3858*) genes were amplified from genomic DNA by PCR with Phusion High-Fidelity DNA Polymerase (NEB) with primers #1155 + #1156 (σ^A^) and #1182 + #1183 (RbpA) and *Msm* chromosomal DNA as the template, cloned into pET22b *via NdeI*/*XhoI* restriction sites and verified by sequencing. Resulting plasmids were transformed into expression *Eco* BL21(DE3) strain resulting in strains LK1740 (σ^A^) and LK1254 (RbpA).

#### HelD

Plasmid encoding the N-terminally His-tagged *Msm* HelD protein was prepared by the *GeneArt*^®^ *Plasmid* Construction Service (Thermofisher). Gene construct for HelD expression was designed by codon-optimized back translation of gene MSMEG_2174 from *Msm* (*strain ATCC 700084 / mc*^*2*^ *155*) with cleavage site for TEV protease placed at the 5’ end. This synthetized gene was cloned into the Champion™ pET302/NT-His expression vector (Thermofisher) *via EcoRI* and *XhoI* restriction sites. Resulting protein thus has 6xHis tag at its N-terminus which is cleavable by TEV protease (protein construct starts with sequence MHHHHHHVNSLEENLYFQG followed by the second amino acid of gene *MSMEG_2174*.

#### HelD-FLAG, σ^A^-FLAG and RbpA-FLAG

The genes coding for the HelD-FLAG, σ^A^-FLAG and RbpA-FLAG proteins were prepared by PCR using Q5^®^ High-Fidelity DNA Polymerase (NEB) with primers #3130 + #3131 (HelD), #2339 + #2340 (σ^A^) and #2894 + #3093 (RbpA) and *Msm* chromosomal DNA as the template. The C-terminal 1x FLAG-tags (DYKDDDDK) were encoded within the reverse PCR primers for all proteins. Subsequently, the genes were inserted into integrative plasmid pTet_INT (PMID: 25711368) via *NdeI*/*HindIII* restriction sites. The constructs were verified by sequencing. Resulting plasmids were transformed into *Msm* mc^2^ 155 (wt, LK865) cells by electroporation resulting in strains LK2590 (HelD-FLAG), LK2073 (σ^A^-FLAG) and LK2541 (RbpA-FLAG).

#### Growth conditions

*Msm* strains - mc^2^ 155 (wt, LK865), σ^A^-FLAG (LK2373), RNAP-FLAG (LK1468), HelD-FLAG (LK2590), and RbpA-FLAG (LK2541) were grown at 37 °C in Middlebrook 7H9 medium with 0.2% glycerol and 0.05% Tween 80 and harvested in exponential phase (OD_600_ ∼0.5; 6 h of cultivation) or early stationary phase (OD_600_ ∼2.5–3.0, 24 h of cultivation) unless stated otherwise. When required, media were supplemented with kanamycin (20 μg/ml). Expression in exponential phase of HelD-FLAG was induced by anhydrotetracycline (1 ng/ml) at 3 h of cultivation. The cells were then grown for additional 3 h. Expressions in stationary phase of σ^A^-FLAG, RbpA-FLAG, and HelD-FLAG were induced by anhydrotetracycline (10 ng/ml) at 8 h of cultivation. The cells were then cultivated for additional 16 h.

### Electron microscopy

#### *Msm* RNAP core purification for cryo-EM

*Eco* strain BL21(DE3) was transformed with pRMS4 (*kanR*) plasmid derivative encoding *Msm* subunits ω, α, and β-β’ fusion with C-terminal His8 tag in one operon from T7 promoter. Expression cultures were incubated at 37 °C and shaken at 250 rpm until OD600 ~0.8, expression was induced with 500 μM isopropyl β-D-thiogalactoside (IPTG) at 17 °C for 16 h. Cells were lysed using sonication by Sonic Dismembrator Model 705 (Fisher Scientific) in a lysis buffer containing 50 mM NaH_2_PO_4_/Na_2_HPO_4_ pH 8 (4 °C), 300 mM NaCl, 2.5 mM MgCl_2_, 30 mM imidazole, 5 mM β-mercaptoethanol, EDTA-free protease inhibitor cocktail (Roche), RNase A (Sigma), DNase I (Sigma) and Lysozyme (Sigma). Clarified lysate was loaded onto a HisTrap FF Crude column (GE Healthcare) and proteins were eluted with a linear gradient of imidazole to the final concentration of 400 mM over 20 column volumes. The *Msm* RNAP core elution fractions were pooled and dialyzed to 20 mM Tris-HCl pH 8 (4 °C), 1 M NaCl, 5% (v/v) glycerol and 4 mM dithiothreitol (DTT) for 20 h. The protein was further polished on XK 26/70 Superose 6 pg column (GE Healthcare) equilibrated in 20 mM Tris-HCl pH 8 (4 °C), 300 mM NaCl, 5% (v/v) glycerol and 4 mM DTT. The *Msm* RNAP core final fractions were eluted at 6 μM concentration, aliquoted, flash-frozen in liquid nitrogen and then stored at –80 °C.

#### *Msm* HelD protein purification for cryo-EM

*Eco* strain Lemo 21 (DE3) was transformed with pET302/NT-His (*cmlR* and *ampR*) plasmid derivative encoding the *Msm* HelD protein fusion with N-terminal 6xHis tag under the control of the T7 promoter. Expression cultures were incubated at 37 °C and shaken at 250 rpm until OD_600_ ~0.8, expression was induced with 500 μM IPTG at 17 °C for 16 h. Cells were lysed using sonication by Sonic Dismembrator Model 705 (Fisher Scientific) in a lysis buffer containing 50 mM Tris-HCl pH 7.5 (4 °C), 400 mM NaCl, 30 mM imidazole, 0.2% Tween20, 2 mM β-mercaptoethanol, EDTA-free protease inhibitor cocktail (Roche), RNase A (Sigma), DNase I (Sigma) and Lysozyme (Sigma). Clarified lysate was loaded onto a HisTrap FF Crude column (GE Healthcare) and proteins were eluted with a linear gradient of imidazole to the final concentration of 400 mM over 20 column volumes. Fractions containing HelD protein were pooled and dialyzed for 20 h against the dialysis buffer containing 20 mM Tris-HCl, pH 7.5 (4 °C), 500 mM NaCl, 1 mM DTT together with TEV protease at a TEV protease:HelD ratio 1:20.

The protein was then concentrated to ~15 A_280_ units and further purified using size-exclusion chromatography using a Superdex 75 column (GE Healthcare) equilibrated in 20 mM Tris-HCl, pH 7.5 (4 °C), 200 mM NaCl and 1 mM DTT. The HelD protein was eluted at ~160 μM concentration, aliquoted, flash-frozen in liquid nitrogen and then stored at –80 °C.

#### *In vitro* HelD-RNAP complex reconstitution for cryo-EM

To assemble the HelD-RNAP complex, the individual proteins were mixed at a molar ratio of 3:1. The *in vitro* reconstitutions were carried out at 4 °C, and the reconstitution mixture was incubated for 15 min. 50 μl of the reconstitution mixture was injected onto a Superose 6 Increase 3.2/300 column (GE Healthcare) equilibrated in 20 mM Tris-HCl, pH 7.8 (4 °C), 150 mM NaCl, 10 mM MgCl_2_ and 1 mM DTT. 50-μl fractions were collected and the protein was eluted at ~1 μM concentration.

#### Electron microscopy

Complexes were diluted to ~850 nM and aliquots of 3 μl were applied to Quantifoil R1.2/1.3 or R2/2 Au 300 mesh grids, immediately blotted for 2 s and plunged into liquid ethane using an FEI Vitrobot IV (4 °C, 100% humidity).

The grids were loaded into an FEI Titan Krios electron microscope at the European Synchrotron Radiation Facility (ESRF) (beamline CM01, ESRF) or CEITEC (Masaryk University, Brno), operated at an accelerating voltage of 300 keV and equipped with a post-GIF K2 Summit direct electron camera (Gatan) operated in counting mode. Cryo-EM data was acquired using EPU software (FEI) at a nominal magnification of x165,000, with a pixel size of 0.8311 and 0.840 Å per pixel. Movies of a total fluence of ~40-50 electrons per Å^2^ were collected at ~1 e^-^/Å^2^ per frame. A total number of 15,177 movies were acquired at a defocus range from −0.7 to −3.3 μm (Table S2).

#### Cryo-EM image processing

All movie frames from three datasets were aligned and dose-weighted using the MotionCor2 program (Figure S3a) and then used for contrast transfer function parameter calculation with Gctf^3^. Initially, particles were selected without a template by Gautomatch (provided by Dr. Kai Zhang, http://www.mrc-lmb.cam.ac.uk/kzhang) from a small portion of the data set (~200 movies). This initial small dataset was subjected to reference-free 2D-classification using RELION 3.0 ^4^. Eight representative classes of different views were selected from the two-dimensional averages and used as reference for automatic particle picking for the dataset I by RELION. WARP^5^ was used for particle picking for datasets II and III.

The resulting particles were iteratively subjected to two rounds of 2D-classification (Figure S3b) at 3x and 2x binned a pixel size. Particles in classes with poor structural features were removed. Particles from dataset I and II were globally refined to estimate the pixel size matching^6^ and particles from dataset II were estimated to match the common pixel size 0.8311 Å per pixel. Particles from all datasets were pooled (~1,560 k), 2x binned and subjected to three-dimensional classifications with image alignment (Figure S4). The first round of 3D-classification was restricted to ten classes and performed using *Msm* RNAP core (PDB entry 6F6W) as a 60 Å low-pass filtered initial model. Classification was done during three rounds of 25 iterations each, using regularization parameter T = 4. During the second and third round, local angular searches were performed at 3.5° and 1.8° to clearly separate structural species. The three most abundant and defined 3D-classes were re-extracted at the pixel size of 0.8311 Å per pixel and 3D auto-refined using respective masks in RELION 3.0 (Figure S4). The results of the 3D auto-refinement were used for per particle CTF refinement in RELION 3.1 ^7^ and further 3D auto-refined. Further 3D classification was applied on class 1 and 3, but no better defined 3D classes were identified. The 3D reconstruction of class 2 was further focus 3D auto-refined on the RNAP core region. The 3D reconstruction of class 2 was also 3D focus classified on the region of the HelD-specific domain and a more defined class was identified and 3D auto-refined separately. The final cryo-EM density maps were generated by the post-processing feature in RELION and sharpened or blurred into MTZ format using CCP-EM^8^. The resolutions of the cryo-EM density maps were estimated at the 0.143 gold standard Fourier Shell Correlation (FSC) cut off (Figure S3e). A local resolution (Figure S3d) was calculated using RELION and reference-based local amplitude scaling was performed by LocScale^9^.

#### Cryo-EM model building and refinement

Atomic models of *Msm* RNAP protein parts (Figure 1a-c) were generated according to the known structure of the *Msm* RNAP core (PDB entry 6F6W). The whole RNAP core was first rigid-body fitted into the cryo-EM density by Molrep^10^ and individual sub-domains fits were optimized using the Jigglefit tool^11^ in Coot^12^ and best fits were chosen according to a correlation coefficient in the JiggleFit tool. The crystal structure of the *Bsu* HelD-2A domain (Figure S7g) was first rigid-body fitted into the cryo-EM density by Molrep^10^ and then manually adapted in Coot. Parts of the HelD main chain were first traced into the cryo-EM density by Buccaneer^13^ and Mainmast^14^.The rest of the HelD protein was built *de-novo* in Coot^11^. The cryo-EM atomic-models of HelD-RNAP complexes were then iteratively improved by manual building in Coot and refinement and validation with Phenix real-space refinement^15^. The atomic models were validated with the Phenix validation tool (Table S2) and the model resolution was estimated at the 0.5 FSC cut-off. Structures were analyzed and Figures were prepared using the following software packages: PyMOL (Schrödinger, Inc.) with APBS plugin^16^, USCF Chimera^17^, CCP4MG^18^, ePISA server^19^.

#### X-ray crystal structure determination of the *Bsu* HelD C-terminal domain

DNA sequence encoding the C-terminal domain of HelD (from residue 608 to 774) was amplified by PCR and cloned into pET15b vector by *NdeI* and *BamHI* restriction sites to make an N-terminal His6-tagged protein. Bacterial culture containing BL21(DE3) RIPL codon-plus cells transformed with a pET15b–HelD-CTD vector was grown at 37 °C in LB medium supplemented with 100 μg/ml ampicillin, protein expression was induced with 0.5 mM IPTG at OD_600_ = 0.5, and incubated for additional 3 h to allow protein expression. Cells were harvested by centrifugation and lysed by sonication in lysis buffer (50 mM Tris-HCl, pH 8.0 at 4 °C, 200 mM NaCl, 5% glycerol, 2 mM β-mercaptoethanol, 2 mM phenylmethylsulfonyl fluoride, PMSF). The lysate was clarified by centrifugation and HelD-CTD was purified by Ni-NTA, Q-sepharose and Heparin-column chromatography. Fractions containing HelD-CTD were concentrated using VivaSpin concentrators until 10 mg/ml in crystallization buffer (10 mM Tris-HCl, pH 8 at 4 °C, 50 mM NaCl, 1 % glycerol, 0.1 mM EDTA, 1 mM DTT).

Crystallization condition of HelD-CTD was screened by using JCSG+ screen (Molecular Dimensions) and crystals were obtained in crystallization solution (0.1 M Na/K phosphate, pH 6.2, 0.2 M NaCl, 50% PEG200) at 22 °C. X-ray crystallographic data were collected at the Penn State X-ray Crystallography Facility and the data were processed by HKL2000^20^. For Sulfur single-wavelength anomalous dispersion phasing, 10 S atom positions were identified and the initial phase and density-modified map were calculated by AutoSol followed by automated model building by AutoBuild in the program Phenix^15^. Iterative refinement by Phenix and model building using Coot^12^ improved the map and model. Finally, water molecules were added to the model. The data statistics and X-ray structure parameters are shown in Table S3.

### Biochemical assays

#### Protein purification for biochemical assays

##### *Msm* RNAP core

Strain of *Eco* containing plasmid with subunits of the RNAP core (LK1853^1^) was grown to the exponential phase (OD_600_ ~ 0.5). Expression of RNAP was induced with 500 μM IPTG for 4 h at room temperature. Cells were harvested by centrifugation, washed, resuspended in P buffer (300 mM NaCl, 50 mM Na_2_HPO_4_, 5% glycerol, 3 mM β-mercaptoethanol) and disrupted by sonication. Cell debris was removed by centrifugation and supernatant was mixed with 1 ml Ni-NTA Agarose (Qiagen) and incubated for 90 minutes at 4 °C with gentle shaking. Ni-NTA Agarose with bound RNAP was loaded on a Poly-Prep^®^ Chromatography Column (BIO-RAD), washed with P buffer and, subsequently, washed with P buffer with 30 mM imidazole. The proteins were eluted with P buffer containing 400 mM imidazole and fractions containing RNAP were pooled and dialysed against storage buffer (50 mM Tris–HCl, pH 8.0, 100 mM NaCl, 50% glycerol, 3 mM β-mercaptoethanol). The RNAP protein was stored at –20 °C.

##### *Msm* σ^A^

Expression strain of *Eco* containing plasmid with gene of σ^A^ (LK1740, this work) was grown at 37 °C until OD_600_ reached ~0.5; expression of σ^*A*^ was induced with 300 μM IPTG at room temperature for 3 h. Isolation of σ^A^ was done in the same way as RNAP purification with the exception of 50 mM imidazole added to the P buffer before resuspending the cells. Instead of the purification in a column, batch purification and centrifugation were used to separate the matrix and the eluate.

##### *Msm* RbpA

The expression and purification of RbpA (LK1254, this work) were done in the same way as for RNAP except when OD_600_ reached ~0.5, the expression was induced with 800 μM IPTG at room temperature for 3 h.

##### *Msm* HelD

*Msm* HelD was prepared as described previously, in the paragraph about purification of proteins for cryo-EM experiments.

Purity of all purified proteins was checked by SDS-PAGE gel.

#### *Msm* HelD ATP, GTP and CTP hydrolysis assay

Hydrolysis of ATP, GTP and CTP (Sigma-Aldrich) by *Msm* HelD was measured in a total volume of 50 μl reaction mixture which contained 10 mM substrate, 10 μg of *Msm* HelD and reaction buffer composed of 50 mM Tris-HCl, pH 7.5, 50 mM NaCl, 5 mM MgCl_2_. Incubation was carried out at 37 °C for 30 min. The amount of released phosphate was analyzed spectrophotometrically at λ = 850 nm according to a modified molybdenum blue method^21^ using a microplate reader Clariostar (BMG LABTECH, Ortenberg, Germany). Briefly, the reaction was stopped by adding 62 μl of reagent A (0.1 M L-ascorbic acid, 0.5 M Cl_3_CCOOH). After thorough mixing, 12.5 μl of reagent B (10 mM (NH_4_)_6_Mo_7_O_24_) and 32 μl of reagent C (0.1 M sodium citrate, 0.2 M NaAsO_2_, 10% acetic acid) was added. All enzymatic reactions were performed in triplicates with separate background readings for each condition.

#### DNA-Protein interaction analysis *in vitro*

DNA-Protein interactions were analyzed on 4-16% Bis-Tris native gels (Thermo Fisher Scientific, cat. No. BN1002BOX) by Electrophoretic Mobility Shift Assay (EMSA). DNA fragment was amplified by Expand High Fidelity PCR System (Roche, cat. No. 11732650001) using #1101 and #1146 primers and *Msm* chromosomal DNA. The resulting 304 bp long PCR fragment was excised and purified from agarose gel. Binding reactions were performed in 1xSTB buffer (50 mM Tris-HCl pH 8.0; 5 mM Mg(C_2_H_3_O_2_)_2_; 100 μM DTT; 50 mM KCl; 50 μg/ml BSA) that contained RNAP (25 pmol), HelD (125 pmol) and DNA (0.2 pmol). First, RNAP was pre-incubated in the presence or absence of HelD (at 37 °C, 45 min). Subsequently, DNA was added and samples were incubated at 37 °C for additional 45 min. Then, NativePage buffer (Invitrogen, cat. No. BN2003) was added and samples were loaded on native gel. Electrophoresis was run in cold room (4 °C). Finally, the gel was stained with DNA stain GelRed nucleic acid stain (Biotium, cat. No. 41003) in 1xTBS for 25 minutes and images were taken with an Ingenius UV-light camera (Syngen). Unbound DNA was quantified by the Quantity One software (BioRad). The gel was subsequently stained with Simply Blue (Invitrogen, cat. No. LC6060) for protein visualization.

#### Protein-Protein interaction analysis *in vitro*

Protein-Protein interactions were analyzed on 7% Tris-acetate native gels (Thermo Fisher Scientific, cat. No. EA0355BOX) by EMSA. Binding reaction was done in 20 μl of 1xSTB buffer containing RNAP (25 pmol), HelD (125 pmol), σ^A^ (1,250 pmol) and RbpA (1,250 pmol) - protein combinations in reactions are specified in the Figure 4 legend. First, RNAP was reconstituted with/without HelD (at 37 °C, 45 min). Then RbpA and/or σ^A^ were added, followed by additional incubation at 37 °C for 45 min. 20 μl of Native Tris-Glycine buffer (Invitrogen, cat. No. LC2673) was added and 20 μl of the mixture was then loaded on a native gel. Electrophoresis was run in cold room (4 °C). Subsequently, for protein visualization, the gels were stained with Simply Blue. The identity of proteins in each band was determined by MALDI mass spectrometric identification.

#### Disassembly of elongation complexes

Elongation complexes (ECs), containing a transcription bubble, were assembled with the *Msm* RNAP core as described before^22^. DNA and RNA oligonucleotides were purchased and are the same as in Table EV7 in ^23^. The RNA (LK-pRNA) was monophosphorylated at the 5’ end by the manufacturer. A 2-fold molar excess of RNA was mixed with template DNA (LK632) in water and annealed in a cycler (45 °C for 2 min, 42-27 °C: temperature was decreasing by 3 °C every 2 min, 25 °C for 10 min). RNAP (32 pmol per sample) was incubated with 4 pmol of the annealed hybrid in 10 μl of reaction buffer (40 mM Tris-HCl, pH 8.0, 10 mM MgCl_2_, 1 mM DTT) for 15 min at room temperature with gentle shaking. 8 pmol of non-template DNA (LK631) containing biotin at the 5’ end was added and the mixture was incubated at 37 °C for 10 min.

Streptavidin-coated magnetic beads (25 μl per sample; Sigma S-2415) were washed with 500 μl of binding buffer (20 mM Tris-CHl, pH 8.0, 0.15 M NaCl) and resuspended in the same volume of fresh binding buffer. Assembled elongation complexes were then mixed with washed beads. ECs and beads were incubated together for 30 min at RT (room temperature) with continuous gentle shaking. Unbound complexes were removed by subsequent washing with 500 μl of binding buffer, 500 μl of washing buffer (20 mM Tris-HCl pH 8.0, 0.5 M NaCl, 2 mM MgCl_2_, 1 mM DTT) and 500μl of reaction buffer^24^. Beads were resuspended in reaction buffer with 100 mM final concentration of KCl, with or without GTP or ATP (final concentration 200 μM) in a total volume of 5 μl. HelD in 2-fold ratio over RNAP (64 pmol per sample) or heat-inactivated HelD (5 min at 95 °C) or buffer were added to the final reaction volume of 10 μl. Reactions proceeded for 20 min at 37 °C. The bound (in complex with EC) and released (free in buffer) RNAPs were separated by using a DYNAL Invitrogen bead separation device. Subsequently, the fractions containing released RNAPs were spotted directly on nitrocellulose membrane. RNAPs were detected by Western blotting using mouse monoclonal antibodies against the β subunit of RNAP (clone name 8RB13) and secondary antibodies conjugated with a fluorophore dye (WesternBrightTM MCF-IR, Advansta, 800 nm anti-mouse antibody) and scanned with an Odyssey reader (LI-COR Biosciences). The analysis was done with the Quantity One software (BioRad). The experiment was conducted in five biological replicates.

#### Immunoprecipitation

150 ml of *Msm* exponential (Figure S9) and 100 ml of stationary phase (Figures 4f, S1, S10) cells were pelleted and resuspended in 4 ml of Lysis buffer (20 mM Tris-HCl, pH 8, 150 mM KCl, 1 mM MgCl_2_) with 1 mM DTT, 0.5 mM PMSF and Sigma protease inhibitor cocktail P8849 (5 μl/ml), sonicated 15 × 10 s with 1 min pauses on ice and centrifuged. 1 ml of stationary and 1.5 ml of exponential phase cells lysates were incubated over night at 4 °C with 25 μl of ANTI-FLAG^®^ M2 Affinity Agarose Gel (Sigma, A2220). Agarose gel beads with the captured protein complexes were washed 4x with 0.5 ml 20 mM Tris-HCl, pH 8, 150 mM KCl, 1 mM MgCl_2_. FLAG-tagged proteins were eluted by 60 μl of 3x FLAG^®^ Peptide (Sigma F4799) (diluted in Tris-buffered saline (TBS) to a final concentration of 150 ng/ml). Proteins were resolved on sodium dodecylsulphate-polyacrylamide gel electrophoresis (SDS-PAGE) and Simply Blue-stained (SimplyBlue, Invitrogen) or analyzed by Western blotting.

#### Double pull-down

Eluted proteins from the first immunoprecipitation (ANTI-FLAG, see above) from lysates of the HelD-FLAG culture from exponential phase were incubated (O/N, 4°C) with 5 μg of σ^A^ or IgG antibodies (negative control), respectively, bound to 20 μl of Protein G-plus Agarose (Santa Cruz Biotechnology, Cat. No. sc-2002), and then 4x washed with 1 ml Lysis buffer. Finally, proteins were analyzed by SDS-PAGE and Western blot.

#### Western blotting

Proteins were resolved by SDS-PAGE and detected by Western blotting using mouse monoclonal antibodies against σ^70^/σ^A^ (clone name 2G10, Biolegend, cat. No. 663208), against the β subunit of RNAP (clone name 8RB13, Biolegend, cat. No. 663903), monoclonal ANTI-FLAG (clone M2, Sigma cat. No. F1814), and anti-mouse secondary antibodies conjugated with HRP (Sigma, cat. No. A7058). Subsequently, the blot was incubated for 5 min with SuperSignal™ West Pico PLUS Chemiluminiscent substrate (Thermo scientific, cat. No. 34577), exposed on film and developed.

#### Trypsin digestion and MALDI mass spectrometric identification

Simply Blue-stained protein bands were cut out from gels, chopped into small pieces and destained using 50 mM 4-ethylmorpholine acetate (pH 8.1) in 50% acetonitrile (MeCN). The gel pieces were then washed with water, reduced in size by dehydration in MeCN and partly dried in a SpeedVac concentrator. The proteins were digested overnight at 37 °C using sequencing grade trypsin (100 ng; Promega) in a buffer containing 25 mM 4-ethylmorpholine acetate and 5% MeCN. The resulting peptides were extracted with 40% MeCN/0.2% TFA (trifluoroacetic acid).

For MALDI MS analysis, 0.5 μl of each peptide mixture was deposited on the MALDI plate, air-dried at room temperature, and overlaid with 0.5 μl of the matrix solution (α-cyano-4-hydroxycinnamic acid in 50% acetonitrile/0.1% TFA; 5 mg/ml, Sigma). Peptide mass maps of proteins in Figures 4f, S1 were measured using an Autoflex Speed MALDI-TOF instrument (Bruker Daltonics, Billerica, USA) in a mass range of 700-4000 Da and calibrated externally using a PepMix II standard (Bruker Daltonics). For protein identification, MS spectra were searched against NCBIprot_20190611 database subset of bacterial proteins using the in-house MASCOT v.2.6 search engine with the following settings: peptide tolerance of 20 ppm, missed cleavage site set to one and variable oxidation of methionine. The spectra of proteins in Figure 4h were acquired on a 15T Solarix XR FT-ICR mass spectrometer (Bruker Daltonics) in a mass range of 500-6000 Da and calibrated internally using peptide masses of *Msm* RpoB and RpoC proteins. The peak lists generated using DataAnalysis 5.0 program were searched against UniProtKB database of *Msm* proteins using the in-house MASCOT engine with the following settings: peptide tolerance of 3 ppm, missed cleavage site set to two and variable and oxidation of methionine.

## Acknowledgments

We thank the ESRF (especially Michael Hons), IBS and EMBL for access to the ESRF Krios beamline CM01; the CEITEC and the Czech Infrastructure for Integrative Structural Biology (CIISB) for access to the CEITEC Krios microscope and to the CMS facilities at BIOCEV (project LM2015043 by MEYS). This work was supported by 20-12109S (to LK) and 20-07473S (to JH) from the Czech Science Foundation, NIH grant R35 GM131860 to KM, and by the Academy of Sciences of the Czech Republic (RVO: 86652036), MEYS (CZ.1.05/1.1.00/02.0109), European Regional Development Fund (Project CIISB4HEALTH, No. CZ.02.1.01/0.0/0.0/16_013/0001776 and ELIBIO, No. CZ.02.1.01/0.0/0.0/15_003/0000447), T Kou. holds a fellowship from the EMBL Interdisciplinary Postdocs (EI3POD) initiative co-funded by Marie Skłodowska-Curie grant agreement No. 664726.

## Author contributions

JD and LK conceived and supervised the project. TKou, TKov, MT, and JDu expressed and purified proteins for cryo-EM, TKou prepared cryoEM grids, collected cryoEM data together with JN, performed image processing and 3D reconstruction, and built initial models together with JDo. KSM and UC solved the *Bsu* HelD CTD. MJ, JH, BB, MŠ, JP, PS, and HŠ did cloning, protein purifications and IPs. JP and PS performed DNA binding experiments. MT, JDu, and TKov performed NTP hydrolysis experiments. TKou, TKov and JDo built and refined atomic models and created figures. IB and MS performed *in silico* modelling. TKou, TKov, JDo and LK wrote the manuscript with input from IB. TS performed initial modelling and comparative analysis.

## Data Availability

Co-ordinates and structure factors or maps have been deposited in the wwwPDB or EMDB.

*Bsu* HelD C-terminal domain (X-ray) PDB ID 6VSX

*Msm* HelD-RNAP complex State I (cryoEM) EMD-10996, PDB ID 6YXU

*Msm* HelD-RNAP complex State II (cryoEM) EMD-11004, PDB ID 6YYS

*Msm* HelD-RNAP complex State III (cryoEM) EMD-11026, PDB ID 6Z11

**Movie 1**: Cryo-EM structure of *Msm* HelD-RNAP complexes in State I – PCh-engaged engaged

**Movie 2:** Cryo-EM structure of *Msm* HelD-RNAP complexes in State II – PCh-engaged and AS-interfering

**Movie 3:** Cryo-EM structure of *Msm* HelD-RNAP complexes in State III – PCh dis-engaged and AS-interfering

## References

1 Kouba, T. et al. The Core and Holoenzyme Forms of RNA Polymerase from Mycobacterium smegmatis. J Bacteriol 201, doi:10.1128/JB.00583-18 (2019).

2 Paget, M. S. Bacterial Sigma Factors and Anti-Sigma Factors: Structure, Function and Distribution. Biomolecules 5, 1245–1265, doi:10.3390/biom5031245 (2015).

3 Lopez de Saro, F. J., Yoshikawa, N. & Helmann, J. D. Expression, abundance, and RNA polymerase binding properties of the delta factor of Bacillus subtilis. J Biol Chem 274, 15953–15958, doi:10.1074/jbc.274.22.15953 (1999).

4 Keller, A. N. et al. epsilon, a new subunit of RNA polymerase found in gram-positive bacteria. J Bacteriol 196, 3622–3632, doi:10.1128/JB.02020-14 (2014).

5 Delumeau, O. et al. The dynamic protein partnership of RNA polymerase in Bacillus subtilis. Proteomics 11, 2992–3001, doi:10.1002/pmic.201000790 (2011).

6 Jensen, D., Manzano, A. R., Rammohan, J., Stallings, C. L. & Galburt, E. A. CarD and RbpA modify the kinetics of initial transcription and slow promoter escape of the Mycobacterium tuberculosis RNA polymerase. Nucleic Acids Res 47, 6685–6698, doi:10.1093/nar/gkz449 (2019).

7 Fairman-Williams, M. E., Guenther, U. P. & Jankowsky, E. SF1 and SF2 helicases: family matters. Curr Opin Struct Biol 20, 313–324, doi:10.1016/j.sbi.2010.03.011 (2010).

8 Wiedermannova, J. et al. Characterization of HelD, an interacting partner of RNA polymerase from Bacillus subtilis. Nucleic Acids Res 42, 5151–5163, doi:10.1093/nar/gku113 (2014).

9 Koval, T. et al. Domain structure of HelD, an interaction partner of Bacillus subtilis RNA polymerase. FEBS Lett 593, 996–1005, doi:10.1002/1873-3468.13385 (2019).

10 Meeske, A. J. et al. High-Throughput Genetic Screens Identify a Large and Diverse Collection of New Sporulation Genes in Bacillus subtilis. PLoS Biol 14, e1002341, doi:10.1371/journal.pbio.1002341 (2016).

11 Krissinel, E. & Henrick, K. Inference of macromolecular assemblies from crystalline state. J Mol Biol 372, 774–797, doi:10.1016/j.jmb.2007.05.022 (2007).

12 Ross, W. et al. ppGpp Binding to a Site at the RNAP-DksA Interface Accounts for Its Dramatic Effects on Transcription Initiation during the Stringent Response. Mol Cell 62, 811–823, doi:10.1016/j.molcel.2016.04.029 (2016).

13 Abdelkareem, M. et al. Structural Basis of Transcription: RNA Polymerase Backtracking and Its Reactivation. Mol Cell 75, 298–309 e294, doi:10.1016/j.molcel.2019.04.029 (2019).

14 Molodtsov, V. et al. Allosteric Effector ppGpp Potentiates the Inhibition of Transcript Initiation by DksA. Mol Cell 69, 828–839 e825, doi:10.1016/j.molcel.2018.01.035 (2018).

15 Laptenko, O., Lee, J., Lomakin, I. & Borukhov, S. Transcript cleavage factors GreA and GreB act as transient catalytic components of RNA polymerase. EMBO J 22, 6322–6334, doi:10.1093/emboj/cdg610 (2003).

16 Perederina, A. et al. Regulation through the secondary channel--structural framework for ppGpp-DksA synergism during transcription. Cell 118, 297–309, doi:10.1016/j.cell.2004.06.030 (2004).

17 Sosunova, E. et al. Donation of catalytic residues to RNA polymerase active center by transcription factor Gre. Proc Natl Acad Sci U S A 100, 15469–15474, doi:10.1073/pnas.2536698100 (2003).

18 Raney, K. D., Byrd, A. K. & Aarattuthodiyil, S. Structure and Mechanisms of SF1 DNA Helicases. Adv Exp Med Biol 973, E1, doi:10.1007/978-1-4614-5037-5_14 (2013).

19 Lee, J. Y. & Yang, W. UvrD helicase unwinds DNA one base pair at a time by a two-part power stroke. Cell 127, 1349–1360, doi:10.1016/j.cell.2006.10.049 (2006).

20 Liu, B., Zuo, Y. & Steitz, T. A. Structural basis for transcription reactivation by RapA. Proc Natl Acad Sci U S A 112, 2006–2010, doi:10.1073/pnas.1417152112 (2015).

21 Tafur, L. et al. Molecular Structures of Transcribing RNA Polymerase I. Mol Cell 64, 1135–1143, doi:10.1016/j.molcel.2016.11.013 (2016).

22 Chen, J. et al. Stepwise Promoter Melting by Bacterial RNA Polymerase. Mol Cell 78, 275–288 e276, doi:10.1016/j.molcel.2020.02.017 (2020).

23 Lin, W. et al. Structural Basis of Transcription Inhibition by Fidaxomicin (Lipiarmycin A3). Mol Cell 70, 60–71 e15, doi:10.1016/j.molcel.2018.02.026 (2018).

24 Boyaci, H. et al. Fidaxomicin jams Mycobacterium tuberculosis RNA polymerase motions needed for initiation via RbpA contacts. Elife 7, doi:10.7554/eLife.34823 (2018).

25 Kohler, R., Mooney, R. A., Mills, D. J., Landick, R. & Cramer, P. Architecture of a transcribing-translating expressome. Science 356, 194–197, doi:10.1126/science.aal3059 (2017).

26 Pani, B. & Nudler, E. Mechanistic insights into transcription coupled DNA repair. DNA Repair (Amst) 56, 42–50, doi:10.1016/j.dnarep.2017.06.006 (2017).

27 Lang, K. S. & Merrikh, H. The Clash of Macromolecular Titans: Replication-Transcription Conflicts in Bacteria. Annu Rev Microbiol 72, 71–88, doi:10.1146/annurev-micro-090817-062514 (2018).

28 Harden, T. T. et al. Alternative transcription cycle for bacterial RNA polymerase. Nat Commun 11, 448, doi:10.1038/s41467-019-14208-9 (2020).

29 Tran, Q. H. & Unden, G. Changes in the proton potential and the cellular energetics of Escherichia coli during growth by aerobic and anaerobic respiration or by fermentation. Eur J Biochem 251, 538–543, doi:10.1046/j.1432-1327.1998.2510538.x (1998).

30 Chen, J. et al. 6S RNA Mimics B-Form DNA to Regulate Escherichia coli RNA Polymerase. Mol Cell 68, 388–397 e386, doi:10.1016/j.molcel.2017.09.006 (2017).

31 Hnilicova, J. et al. Ms1, a novel sRNA interacting with the RNA polymerase core in mycobacteria. Nucleic Acids Res 42, 11763–11776, doi:10.1093/nar/gku793 (2014).

## Methods references

2 Kim, J. H. et al. Protein inactivation in mycobacteria by controlled proteolysis and its application to deplete the beta subunit of RNA polymerase. Nucleic Acids Res 39, 2210–2220, doi:10.1093/nar/gkq1149 (2011).

3 Zhang, K. Gctf: Real-time CTF determination and correction. J Struct Biol 193, 1–12, doi:10.1016/j.jsb.2015.11.003 (2016).

4 Zivanov, J. et al. New tools for automated high-resolution cryo-EM structure determination in RELION-3. Elife 7, doi:10.7554/eLife.42166 (2018).

5 Tegunov, D. & Cramer, P. Real-time cryo-electron microscopy data preprocessing with Warp. Nat Methods 16, 1146–1152, doi:10.1038/s41592-019-0580-y (2019).

6 Wilkinson, M. E., Kumar, A. & Casanal, A. Methods for merging data sets in electron cryo-microscopy. Acta Crystallogr D Struct Biol 75, 782–791, doi:10.1107/S2059798319010519 (2019).

7 Zivanov, J., Nakane, T. & Scheres, S. H. W. Estimation of high-order aberrations and anisotropic magnification from cryo-EM data sets in RELION-3.1. IUCrJ 7, 253–267, doi:10.1107/S2052252520000081 (2020).

8 Burnley, T., Palmer, C. M. & Winn, M. Recent developments in the CCP-EM software suite. Acta Crystallogr D Struct Biol 73, 469–477, doi:10.1107/S2059798317007859 (2017).

9 Jakobi, A. J., Wilmanns, M. & Sachse, C. Model-based local density sharpening of cryo-EM maps. Elife 6, doi:10.7554/eLife.27131 (2017).

10 Vagin, A. & Teplyakov, A. Molecular replacement with MOLREP. Acta Crystallogr D Biol Crystallogr 66, 22–25, doi:10.1107/S0907444909042589 (2010).

11 Brown, A. et al. Tools for macromolecular model building and refinement into electron cryo-microscopy reconstructions. Acta Crystallogr D Biol Crystallogr 71, 136–153, doi:10.1107/S1399004714021683 (2015).

12 Emsley, P. & Cowtan, K. Coot: model-building tools for molecular graphics. Acta Crystallogr D Biol Crystallogr 60, 2126–2132, doi:S0907444904019158 [pii], doi 10.1107/S0907444904019158 (2004).

13 Cowtan, K. The Buccaneer software for automated model building. 1. Tracing protein chains. Acta crystallographica. Section D, Biological crystallography 62, 1002–1011, doi:10.1107/S0907444906022116 (2006).

14 Terashi, G. & Kihara, D. De novo main-chain modeling for EM maps using MAINMAST. Nat Commun 9, 1618, doi:10.1038/s41467-018-04053-7 (2018).

15 Afonine, P. V. et al. Real-space refinement in PHENIX for cryo-EM and crystallography. Acta Crystallogr D Struct Biol 74, 531–544, doi:10.1107/S2059798318006551 (2018).

16 Jurrus, E. et al. Improvements to the APBS biomolecular solvation software suite. Protein Sci 27, 112–128, doi:10.1002/pro.3280 (2018).

17 Pettersen, E. F. et al. UCSF Chimera--a visualization system for exploratory research and analysis. J Comput Chem 25, 1605–1612, doi:10.1002/jcc.20084 (2004).

18 McNicholas, S., Potterton, E., Wilson, K. S. & Noble, M. E. Presenting your structures: the CCP4mg molecular-graphics software. Acta Crystallogr D Biol Crystallogr 67, 386–394, doi:10.1107/S0907444911007281 (2011).

19 Schlee, S. et al. Prediction of quaternary structure by analysis of hot spot residues in protein-protein interfaces: the case of anthranilate phosphoribosyltransferases. Proteins 87, 815–825, doi:10.1002/prot.25744 (2019).

20 Otwinowski, Z. & Minor, W. Processing of X-ray diffraction data collected in oscillation mode. Methods Enzymol 276, 307–326 (1997).

21 He, Z. & Honeycutt, C. W. A Modified Molybdenum Blue Method for Orthophosphate Determination Suitable for Investigating Enzymatic Hydrolysis of Organic Phosphates. Communications in Soil Science and Plant Analysis 36, 1373–1383, doi:10.1081/CSS-200056954 (2005).

22 Komissarova, N., Kireeva, M. L., Becker, J., Sidorenkov, I. & Kashlev, M. Engineering of elongation complexes of bacterial and yeast RNA polymerases. Methods Enzymol 371, 233–251, doi:10.1016/S0076-6879(03)71017-9 (2003).

23 Sikova, M. et al. The torpedo effect in Bacillus subtilis: RNase J1 resolves stalled transcription complexes. EMBO J 39, e102500, doi:10.15252/embj.2019102500 (2020).

24 Deng, Z. et al. Yin Yang 1 regulates the transcriptional activity of androgen receptor. Oncogene 28, 3746–3757, doi:10.1038/onc.2009.231 (2009).

