## Supplemental Information for "Mycobacterial HelD is a nucleic acids-clearing factor for RNA polymerase"

This PDF file includes:

Figures S1 to S12

Tables S1 to S3

Supplementary references

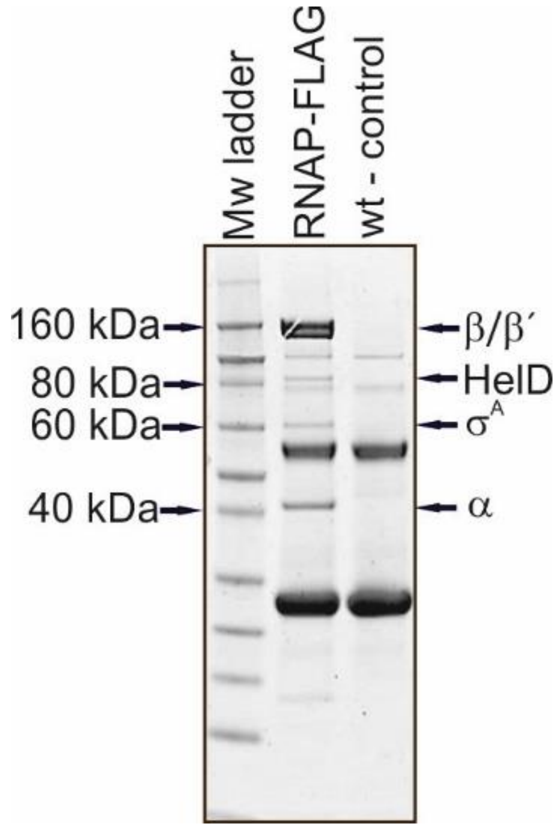


**Figure S1: *Msm* HelD is in complex with RNAP**

SDS-PAGE of IPs of RNAP-FLAG from *Msm* (RNAP-FLAG, strain LK1468; wt – strain LK865). The gel shows boiled ANTI-FLAG M2 agarose with bound proteins. The identities of the pulled-down proteins are indicated with arrows (determined by mass spectrometry). Wt – control, a strain without any FLAG-fusion. The experiment was performed 3x (biological replicates). Mw, molecular weight marker. The two prominent un-marked bands correspond to heavy and light antibody chains, respectively.

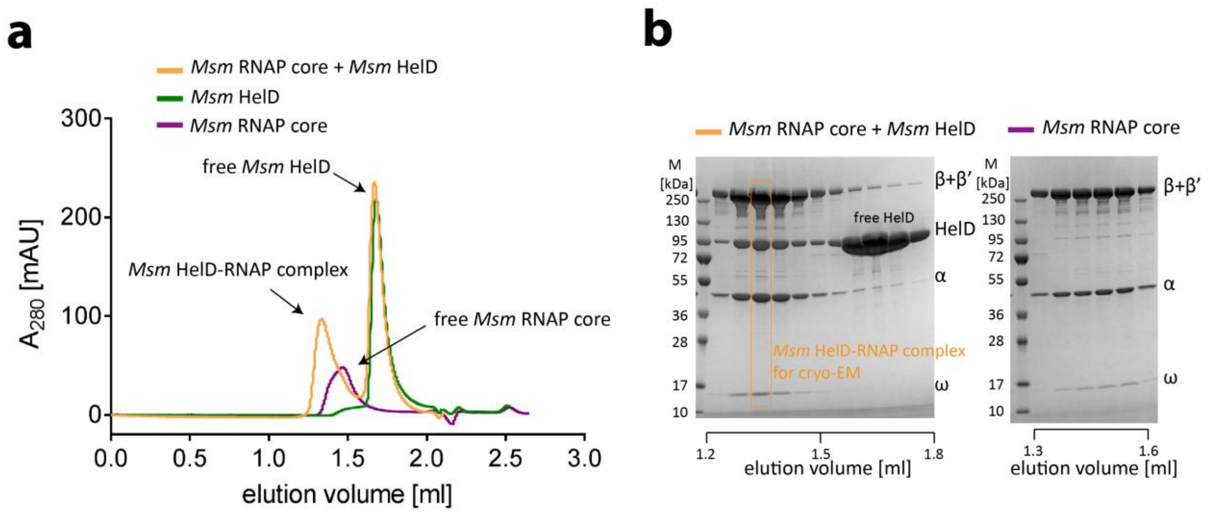


**Figure S2: Reconstitution of *Msm* HelD-RNAP complex.**

**a**, Size-exclusion chromatography (SEC) analysis of RNAP core alone (purple line) and HelD protein alone (green line). SEC analysis of protein sample after reconstitution of RNAP core with HelD protein at a 1:3 ratio (yellow line). The first yellow peak (from left) is the *Msm* HelD-RNAP complex, the second yellow peak is excess of free HelD protein. **b**, SDS-PAGE analysis of the *Msm* HelD-RNAP complex and the *Msm* RNAP core. 40µg protein samples of fractions of *Msm* HelD-RNAP complex and RNAP core alone were loaded onto analytical SDS-PAGE. Fractions are indicated by the elution volume. The first lane contains the molecular weight marker.

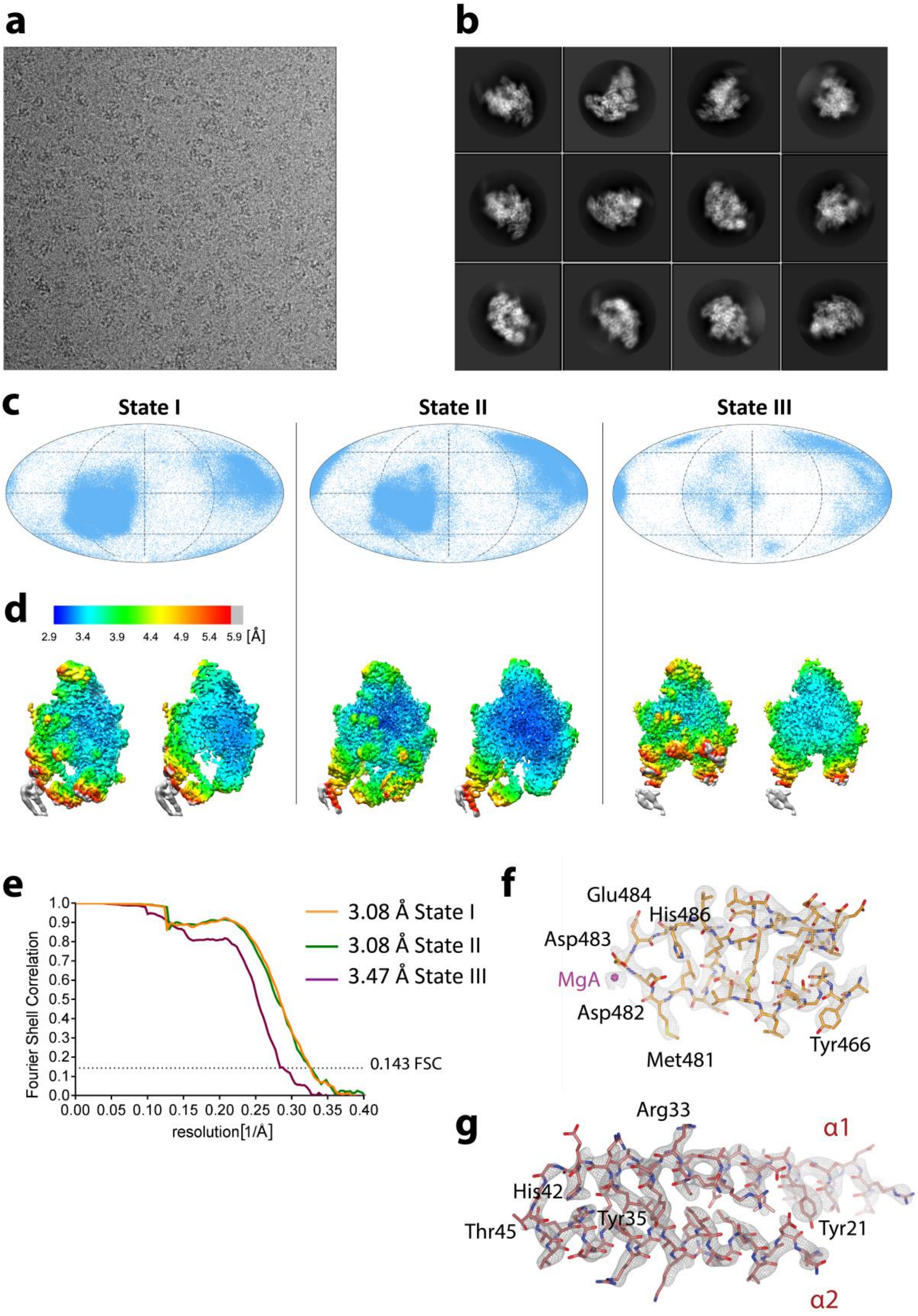
a
b
c
State I
State II
State III
d
2.9 3.4 3.9 4.4 4.9 5.4 5.9 [Å]
e
Fourier Shell Correlation
1.0 0.9 0.8 0.7 0.6 0.5 0.4 0.3 0.2 0.1 0.0
0.00 0.05 0.10 0.15 0.20 0.25 0.30 0.35 0.40
resolution[1/Å]
3.08 Å State I
3.08 Å State II
3.47 Å State III
0.143 FSC
f
Glu484
His486
Asp483
MgA
Asp482
Met481
Tyr466
g
Arg33
α1
His42
Tyr35
Thr45
Tyr21
α2

**Figure S3: Cryogenic electron microscopy of *Msm* HelD-RNAP complex.**

**a**, Representative micrograph of *Msm* HelD-RNAP complex in free standing ice after MotionCor2[1] correction at defocus of ~2.5 µm. **b**, 2D-class averages of the *Msm* HelD-RNAP complex. **c**, Angular distribution for particle projections of the *Msm* HelD-RNAP complex State I, II and III respectively, visualized on a globe-like plane. **d**, Distribution of local resolution of the *Msm* HelD-RNAP State I, II and III, respectively. Surface (**left**) and slice (**right**) representation. Maps are colored according to the local resolution calculated within the RELION software package. Resolution is as indicated in the color bar. **e**, Fourier shell correlation (FSC) curves for *Msm* HelD-RNAP complex State I (yellow), II (green) and III (purple), respectively. The plot of the FSC between two independently refined half-maps shows the overall resolution of the two maps as indicated by the gold standard FSC 0.143 cut-off criteria[2]. **f,** Cryo-EM density for the HelD PCh-loop tip, MgA is shown as magenta sphere. **g,** Cryo-EM density for the N-terminal CC-domain of HelD.

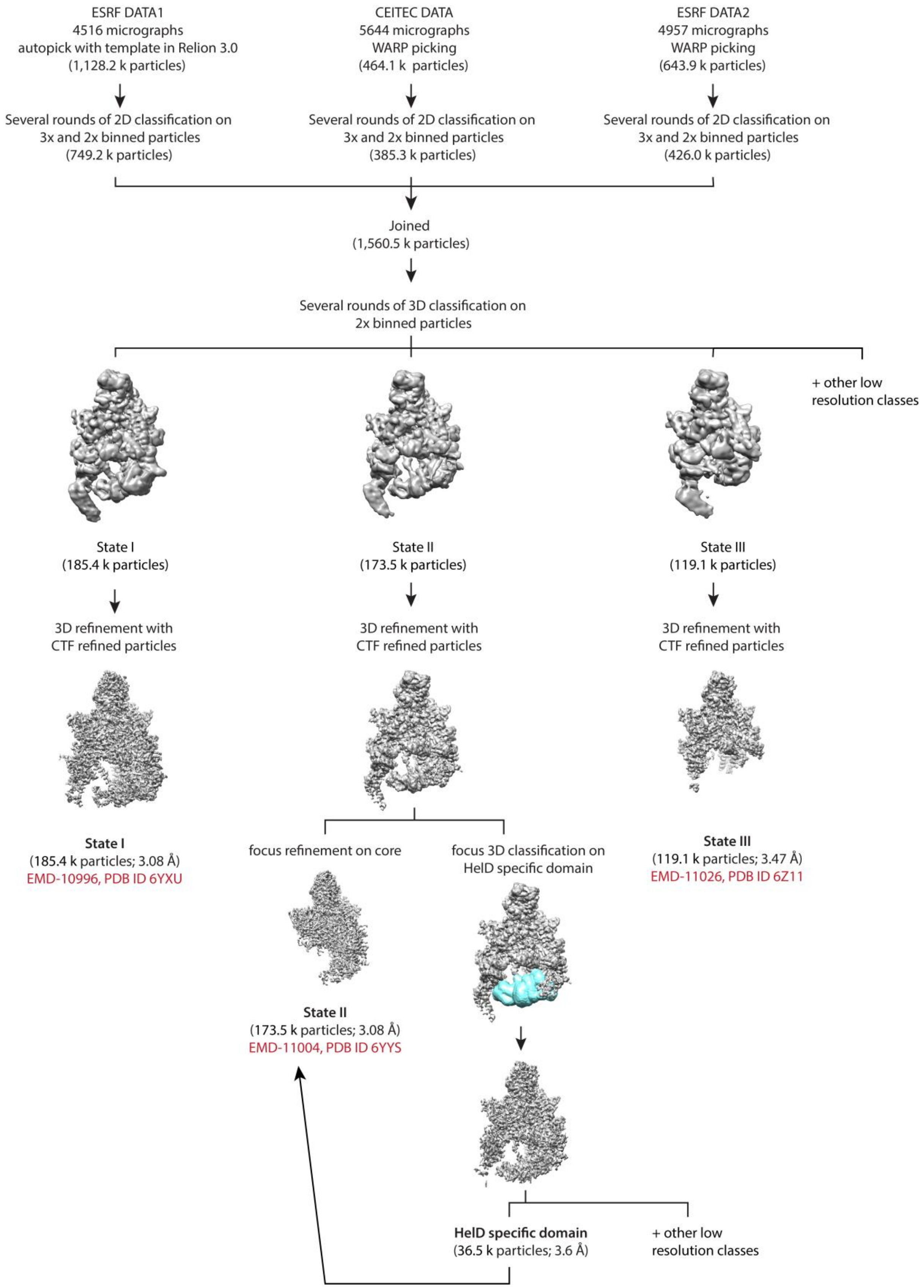
ESRF DATA1
4516 micrographs
autopick with template in Relion 3.0
(1,128.2 k particles)
CEITEC DATA
5644 micrographs
WARP picking
(464.1 k particles)
ESRF DATA2
4957 micrographs
WARP picking
(643.9 k particles)
Several rounds of 2D classification on 3x and 2x binned particles
(749.2 k particles)
Several rounds of 2D classification on 3x and 2x binned particles
(385.3 k particles)
Several rounds of 2D classification on 3x and 2x binned particles
(426.0 k particles)
Joined
(1,560.5 k particles)
Several rounds of 3D classification on 2x binned particles
+ other low resolution classes
State I
(185.4 k particles)
State II
(173.5 k particles)
State III
(119.1 k particles)
3D refinement with CTF refined particles
3D refinement with CTF refined particles
3D refinement with CTF refined particles
State I
(185.4 k particles; 3.08 Å)
EMD-10996, PDB ID 6YXU
focus refinement on core
focus 3D classification on HelD specific domain
State III
(119.1 k particles; 3.47 Å)
EMD-11026, PDB ID 6Z11
State II
(173.5 k particles; 3.08 Å)
EMD-11004, PDB ID 6YYS
HelD specific domain
(36.5 k particles; 3.6 Å)
+ other low resolution classes

**Figure S4: Cryo-EM data 3D classification and refinement scheme.**

Summary of the cryo-EM 3D classification and refinement scheme of the *Msm* HelD-RNAP complex. Initially, three different datasets were processed individually to the level of 2D classification. 2D classes with well-defined secondary structure features were merged (1,560.5k particles). The merged particles were classified into ten 3D classes with angular assignment. Incomplete, low resolution, and damaged particle classes were excluded from further data analyses. The three most prominent 3D classes of the *Msm* HelD-RNAP complex were refined corresponding to State I, II and III. The State II class was focus-refined around the region of the RNAP core and the HelD N-terminal and 1A domain. Another round of focus classification was performed on the region of the HelD 1A and HelD-specific domains using corresponding mask (cyan). Atomic resolution cryo-EM maps were refined and post-processed with their respective masks in RELION 3.0[3,4].

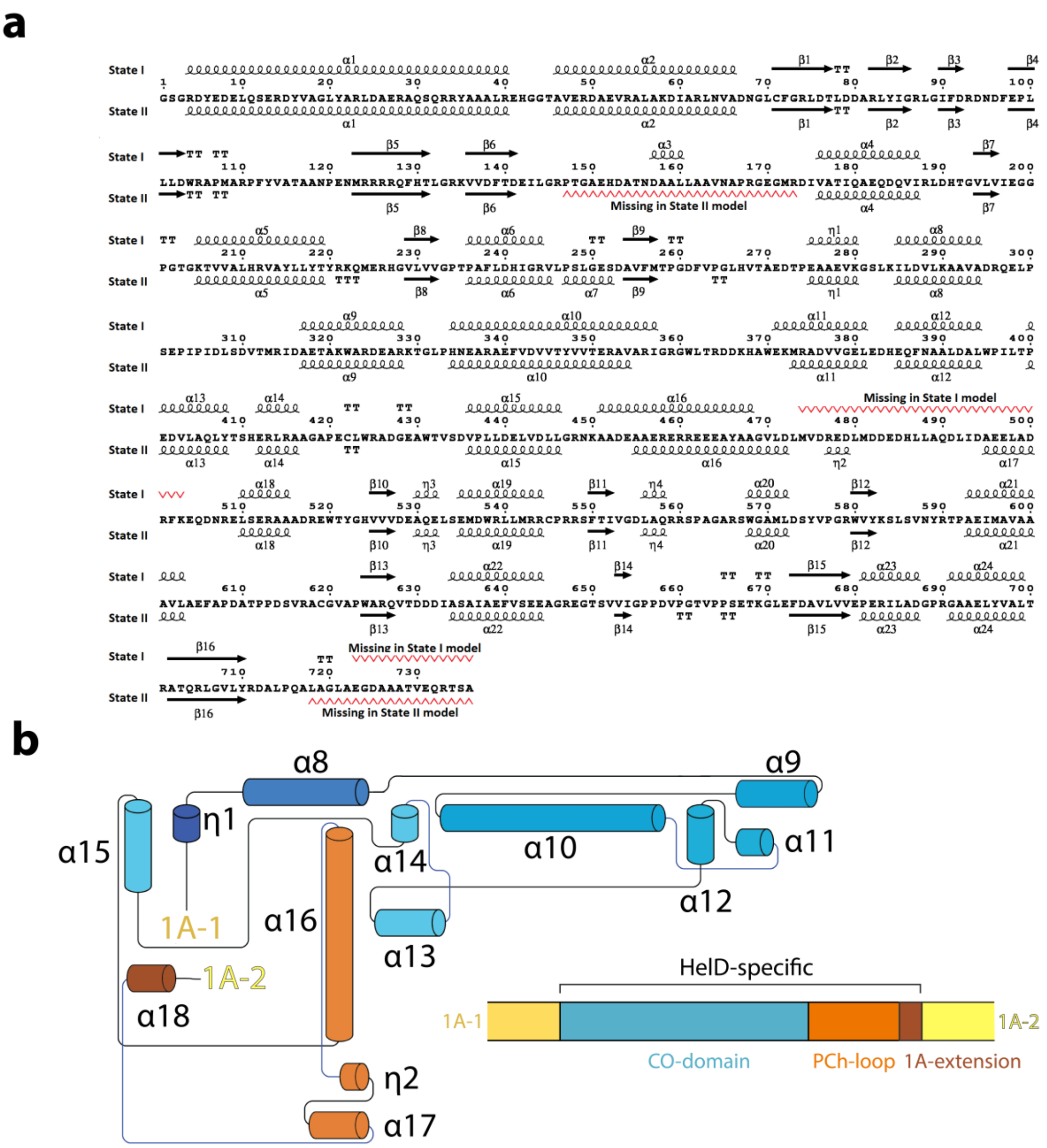


**Figure S5: Secondary structure assignment of HelD protein**

**a**, State I (**top**) and State II (**bottom**) secondary structure elements marked along the *Msm* HelD amino acid sequence. Some regions (red marking) are not folded in one or the other State, α7 exists in State II only, α16 has a shifted register. **b**, Topology of the new fold of the HelD-specific domain (no structural homolog identified).

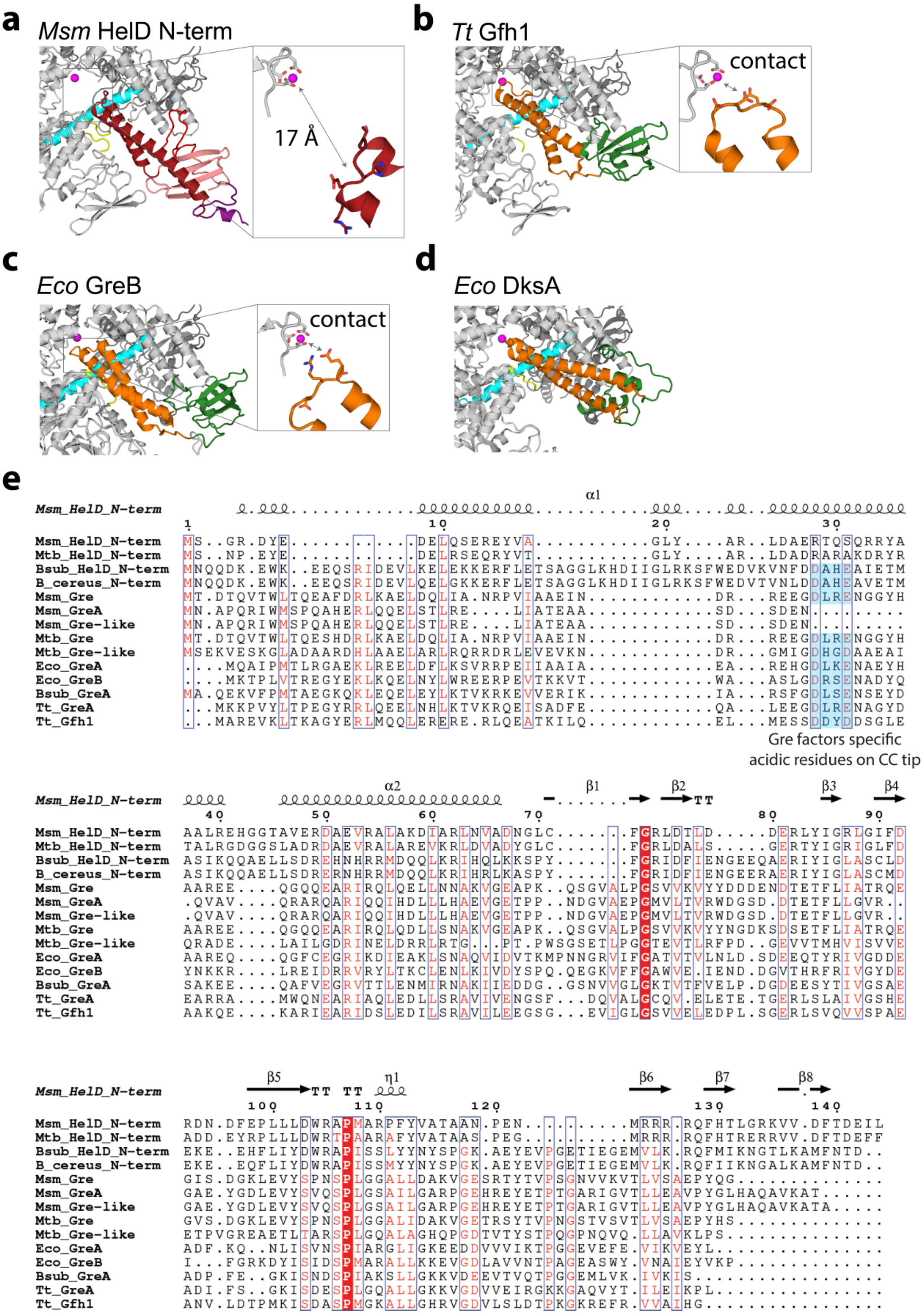
a
Msm HelD N-term
17 Å
b
Tt Gfh1
contact
c
Eco GreB
contact
d
Eco DksA
e
Msm_HelD_N-term
Mtb_HelD_N-term
Bsub_HelD_N-term
B_cereus_N-term
Msm_Gre
Msm_GreA
Msm_Gre-like
Mtb_Gre
Mtb_Gre-like
Eco_GreA
Eco_GreB
Bsub_GreA
Tt_GreA
Tt_Gfh1
Gre factors specific acidic residues on CC tip

**Figure S6: Structural comparison of HelD and Gre-like transcription factors**

**a**, **b**, **c**, and **d** Structural comparison of (**a**) *Msm* HelD N-terminal domain and Gre-like transcription factors. HelD anchors into the RNAP secondary channel similarly to (**b**) *Tt* Gfh1 (PDB ID 3AOH) and (**c**) *Eco* GreB (PDB ID 6RI7) N-terminal CC (orange) and globular (green) domains. However, in contrast to GreB and Gfh1 CC domains, the tip of HelD NCC-domain does not reach to the AS (insets, MgA as magenta sphere). (**d**) *Eco* DksA interacts with the RNAP secondary channel in a similar fashion (PDB ID 5W1T). **e**, Sequence alignment of HelD homologs and Gre-like transcription factors. The mycobacterial HelD NCC-domain tip does not contain the conserved DXX(E/D) motif necessary for Gre factor-like endonuclease activity.

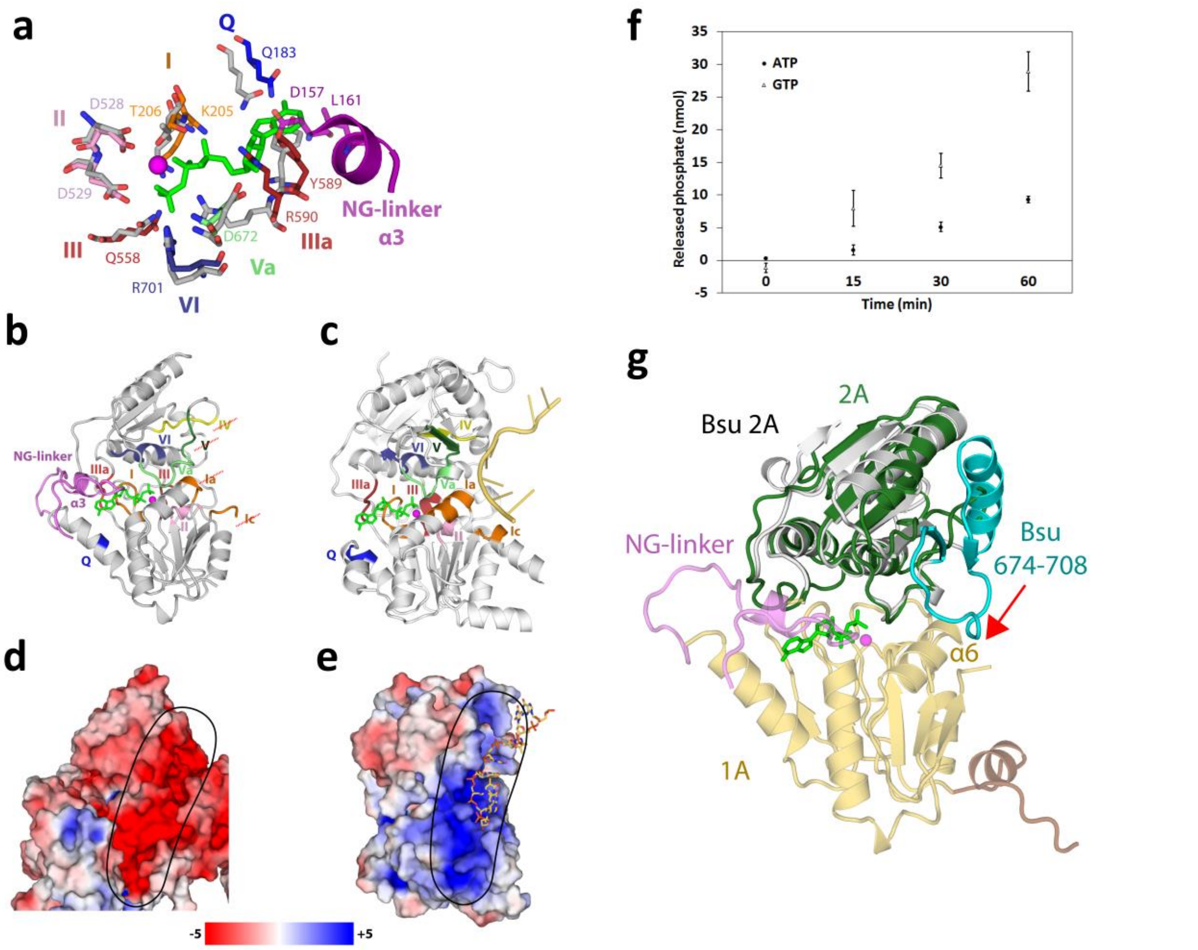


**Figure S7: *Msm* HelD 1A-2A heterodimer nucleotide binding site compared to UvrD; NTPase activity of *Msm* HelD; *Bsu* HelD CTD crystal structure;**

**a**, Superposition of HelD NTP-binding site (State I, color coded) and UvrD ATP-bound state (grey, PDB ID 2IS4[5]). Conserved residues from motifs Q (blue), I (orange), II (pink), ~III and IIIa (firebrick), Va (light green) and VI (deep blue) are present but not in conformations compatible with NTP binding. The ordered NG-linker locks the conformation of Tyr589 (VdW interactions with residues HelD/157, 160 and 161 of α3) and of Arg590 (Arg side chain links Asp157 and Glu672 of HelD) that they would clash with NTP base a ribose, probably making the NTP binding/hydrolysis in State I impossible. **b** and **c**, Conserved nucleotide binding site motifs Q, I, II, III, IIIa, Va, and VI (color coded as in **Figure 2d**) as observed in HelD (**a**,**b**) in comparison to UvrD (**c**,PDB ID 2IS4). Residues responsible for ssDNA (pale yellow in **c**) binding in motifs Ia and Ic (orange), IV (yellow) and V (forest green) in UvrD are not present in HelD (red crosses). **d** and **e**, comparison of surface electrostatic potential of the HelD 1A-2A heterodimer and UvrD ssDNA-bound 1A-2A heterodimer, respectively. A prominent positively charged groove binds ssDNA (sticks in **e**) on the surface of UvrD (black oval). In contrast, a negatively charged groove is present in a similar area of HelD surface (black oval). Electrostatics surfaces were generated by APBS within PyMol according to heat bar in $k_BT/e$ units. **f,** Hydrolyses of ATP and GTP were monitored and evaluated at 0, 15, 30 and 60 min intervals. Measurements were performed in triplicates for each time interval with separate background readings for each condition. The results are shown as amount of released phosphate in the reaction, with standard deviations shown as error bars.

**g**, X-ray structure of the C-terminal domain of *Bsu* HelD compared with State I of *Msm* HelD. The C-terminal domain of *Bsu* HelD (residues 608-773) shown as secondary structure elements in grey superimposed by the SSM algorithm with the 2A domain of *Msm* HelD (colored as in **Figure 1d**); ATP (green sticks) and $Mg^{2+}$ (magenta sphere) in pose as in the structure 2IS4 superimposed according to the NTP-binding site motifs in *Msm* HelD. The 2A domain structure of *Bsu* HelD corresponds to the Rossman fold of the RecA-like domain (central twisted 5-stranded β-sheet surrounded by 5 α-helices 611-620, 645-663, 674-687, 733-745, and 760-764); loop 624-630 was not localized. The domain is most similar to the crystal structure of the C-terminal domain of putative DNA helicase from *Lactobacillus plantarum* (PDB ID 3DMN, rmsd 1.23 Å, 151 aligned $C^{\alpha}$ atoms, 37.7% sequence identity) with identical fold and topology (PDBeFold server[6]). The structure aligns well with that of the 2A/2B domain of UvrD (PDB ID 2IS4, rmsd 1.6 Å, 149 aligned $C^{\alpha}$ atoms), with an almost perfect match of the secondary structure, however of significantly different topology (not shown). The C-terminal domain of *Bsu* HelD has a very similar localization of the amino acid residues forming the expected NTP-binding site (Arg608 corresponds to UvrD/Arg284 – part of motif IIIa, motif VI occurs as 741-TACTRAM-747, Arg745 very likely participating in NTP binding and cleavage, Glu716 is conserved in position of UvrD/Glu566, likely binding the NTP ribose moiety). In comparison with State I of *Msm* HelD the *Bsu* structure is more similar to the 2A domain (rmsd 2.2 Å, 92 aligned residues, sequence identity of the aligned parts 21.7%, alignment shown) than to 1A (2.7 Å, 102 residues aligned, 9.8%, alignment not shown). The helix-loop-strand motif 674-708 (cyan) of *Bsu* HelD does not match any element of 2A in *Msm* HelD and the region 695-699 of the loop would clash with α6 of domain 1A in *Msm* HelD.

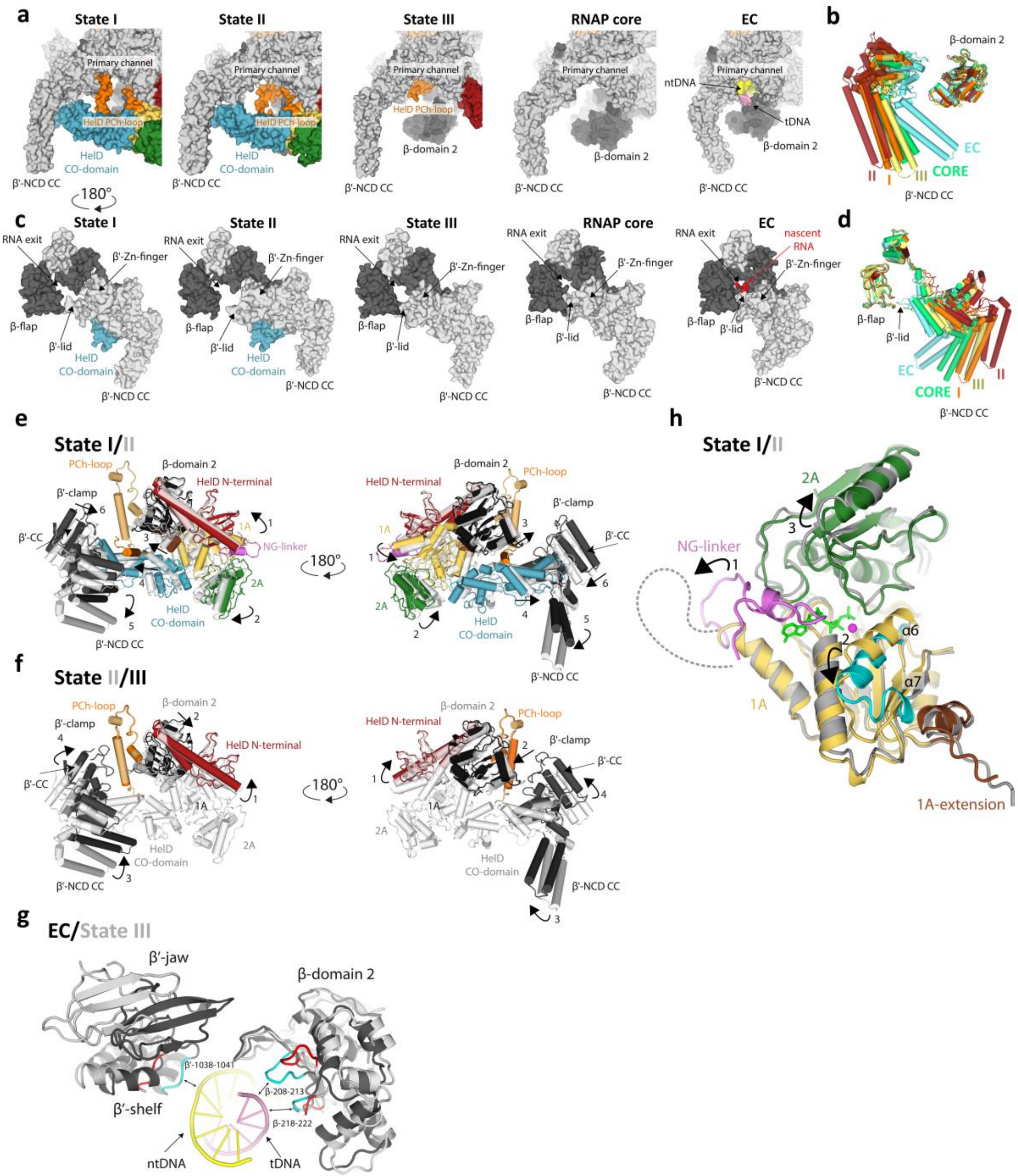


**Figure S8: The *Msm* HelD specific domain wedges into the RNAP primary channel; global domain changes in *Msm* HelD states**

**a**, Surface representation of HelD specific domain interaction with RNAP primary channel in State I, II, and III, compared to *Msm* RNAP core (PDB ID 6F6W) and model of *Msm* elongation complex according to PDB ID 2O5J. Color code as in **Figure 1d**, template DNA in pink, non-template in yellow. **b**, Comparison of RNAP primary channel opening in RNAP complex with HelD in State I (orange), II (red), III (yellow), and without HelD in RNAP core (green) in EC (cyan). **c**, Surface representation of RNA exit channel opening caused by HelD interaction with RNAP in State I, II, and III, compared to *Msm* RNAP core (PDB ID 6F6W) and model of *Msm* elongation complex according to PDB ID 2O5J. Color code as in **Figure 1d**, nascent RNA in red.

**d**, Comparison of RNAP RNA exit channel opening in RNAP complex with HelD in State I (orange), II (red), III (yellow), and without HelD in RNAP core (green) in EC (cyan).

**e,** Two views of State I and II superposition according to the RNAP core (β/430-738). The collapse of NG-linker in State II allows for 1A and 2A mutual reorientation (arrow 1 and 2). Concomitantly this causes a shift of 1A extension (arrow 3 in left panel) and β-domain 2 (arrow 3 in right panel). The reorientation of 1A-2A also causes a shift of the HelD CO-domain (arrow 4) and a further swing-out of β'-NCD CC (arrow 5). On the other hand the β'-CC shifts towards the HelD CO-domain (arrow 6). State I is colored as in **Figure 1**, State II is in light transparent grey. Only selected domains are displayed. **f**, Two views of State II and III superposition according to the RNAP core (β/430-738). In State III, the HelD N-terminal domain slightly shifts within the RNAP secondary channel (arrow 1). The absence of 1A and 2A domains in State III allows relaxation of β-domain 2, which shifts to a similar position as in State I (arrow 2), The absence of the HelD-specific domain allows closure of the β' clamp (arrow 3 and 4). State III is colored as in **Figure 1**, State II is in light transparent grey as in **e**. Only selected domains are displayed. **g**, Superposition of State III (grey) with EC (black) according to the RNAP core (β/430-738), only selected domains are displayed. The HelD N-terminal domain insertion into the secondary channel induces changes in the RNAP primary channel that may destabilise the dwDNA interaction. Notice the shifts of both β-domain 2 and β'-jaw/shelf and changes in the loops (cyan) contacting (double arrows) dwDNA in EC (red) and in the HelD presence. **h**, Superposition of the 1A-2A heterodimer in State I (colored as in **Figure 1**) and State II (light grey) according to 1A-1 domain (1A-1 residues 174-259 superimposed by least squares on main chain, rmsd 2.37 Å). In State II, the disorder of NG-linker (arrow 1), rearrangement of α6 and formation of α7 (change from yellow to cyan, arrow 2), and shift of the 2A domain (arrow 3) altogether result in more open NTP-binding site (ATP in green, $Mg^{2+}$ magenta sphere, modelled by superposition with UvrD ternary complex, PDB ID 2IS4).

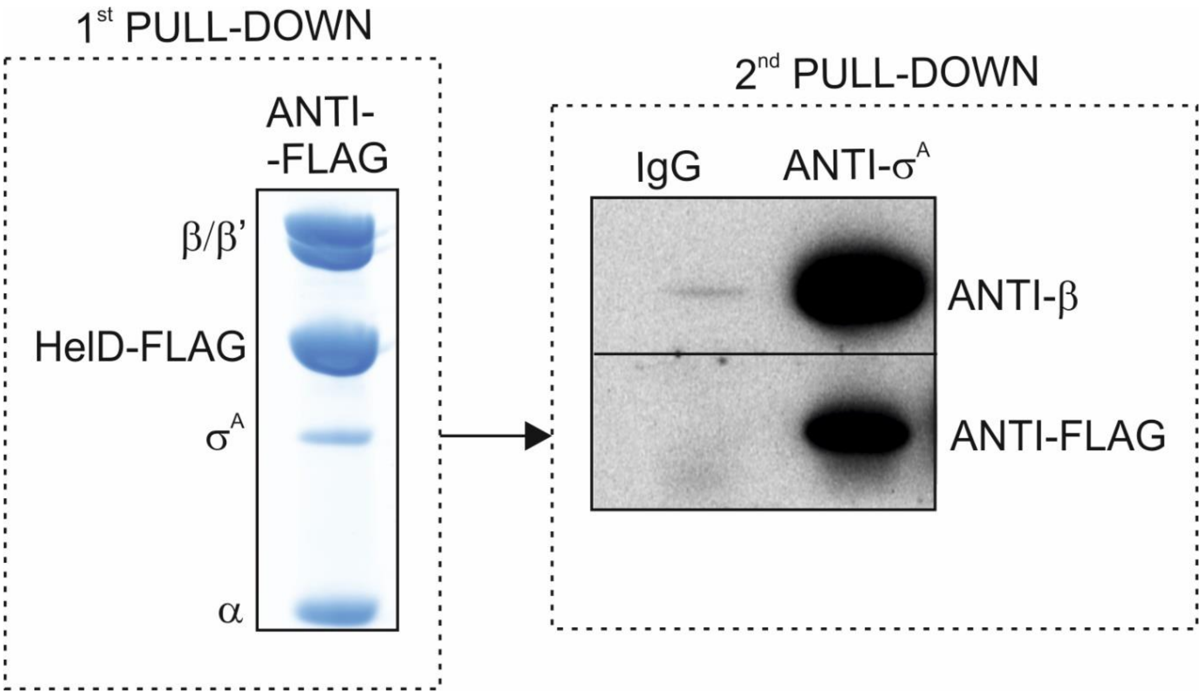


**Figure S9: HelD and $\sigma^A$ can coexist on RNAP.**

Double pull-down: The first pull-down was performed from *Msm* lysates (strain LK2590) with antibody against the FLAG peptide (the same result as in Figure 4f). The Simply Blue-stained gel shows the resulting pulled-down proteins. This protein mixture from the first pull-down was then used for the second pull-down with antibody against $\sigma^A$ and with IgG (negative control). The presence of HelD-FLAG (anti-FLAG) and RNAP (anti-β) was verified by Western blotting. The identities of the antibodies used for the detection are indicated next to the gel. The experiment was performed 2x with identical results.

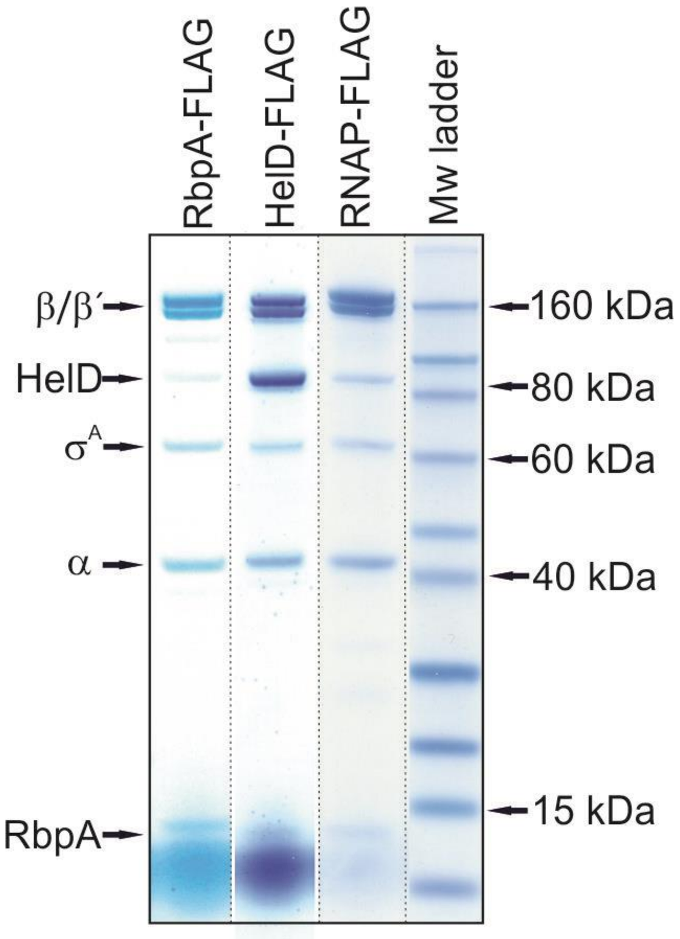


**Figure S10: RbpA is in complex with RNAP, σ^A, and HelD.**

Simply Blue-stained SDS-PAGE of IPs of FLAG-tagged proteins from *Msm* (RbpA-FLAG, strain LK2541; HelD-FLAG, strain LK2590; RNAP-FLAG, strain LK1468). The identities of the FLAG-tagged proteins are indicated above the lanes. The identities of the pulled-down proteins are indicated with arrows (determined by mass spectrometry). The final gel was assembled electronically as indicated with the dotted lines. The experiment was performed 3x (biological replicates).

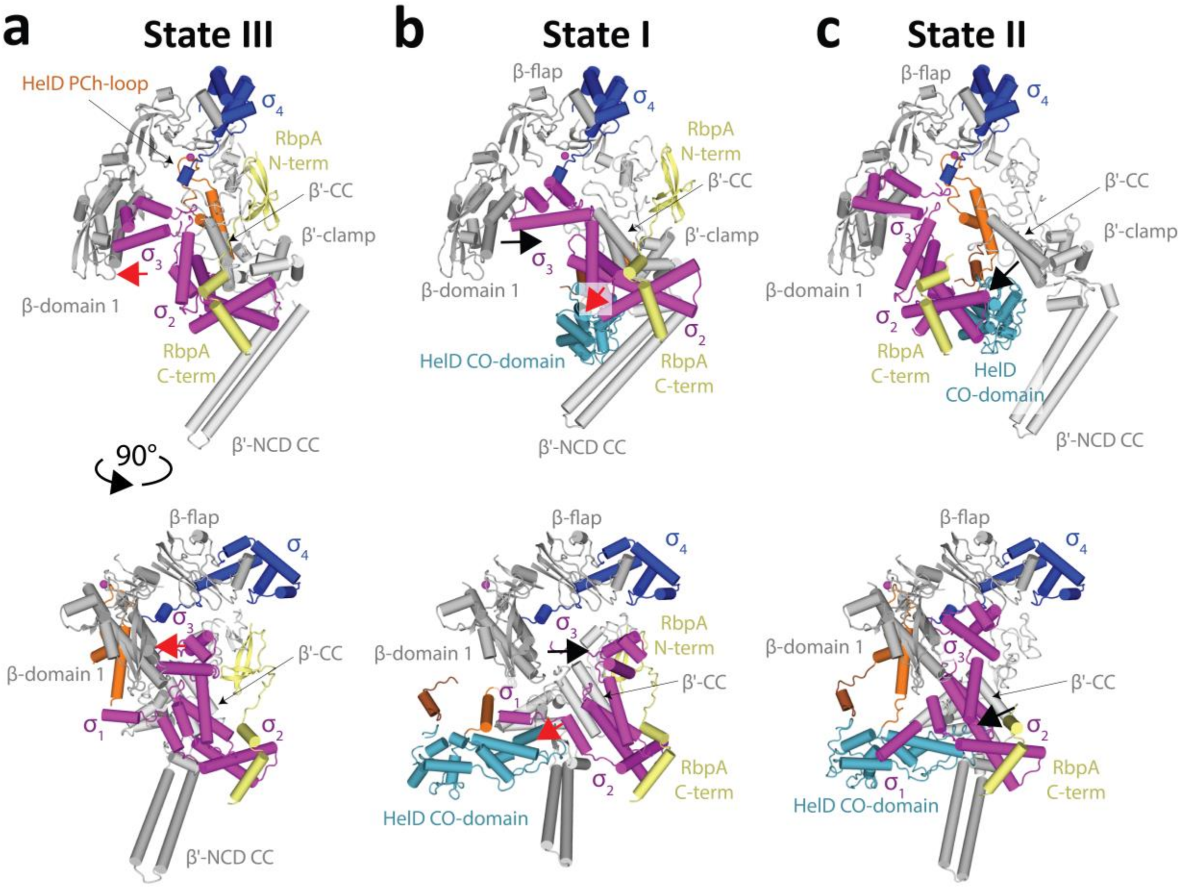


**Figure S11: Models of HelD, $\sigma^A$, and RbpA coexistence on RNAP**

**a**,**b**,**c** Three hypothetical coexistence modes (ordered according to the least adjustments needed) of HelD (only HelD-specific domains shown for clarity), $\sigma^A$, and RbpA in the RNAP primary channel in two perpendicular views. Color code as in **Figure 1**, domains $\sigma^{1\text{-}3}$ magenta, $\sigma^4$ blue, RbpA yellow. **a**, The State III complex superimposed with PDB entries ID 6EYD and ID 5TW1 based on the RNAP core domain (β/430-738). In State III, the HelD CO-domain does not occupy the primary channel, and $\sigma^2$ can interact with the conserved binding site on the β' clamp coiled-coil domain (β'-CC). The $\sigma^3$ domain clashes sterically with β-domain 1 (also called β-protrusion, red arrow), however, a slight shift of $\sigma^3$ could accommodate the latter. The RbpA interaction with both $\sigma^A$ and β' clamp is preserved. **b**, The State I complex superimposed with the PDB entries ID 6EYD and ID 5TW1 based on the RNAP β' clamp (β'/6-404). In State I, the HelD CO-domain occupies the primary channel and $\sigma^2$ can interact with β'-CC if the CO-tip accommodates for $\sigma^2$ presence (red arrow) and $\sigma^1$ moves away. The opening of the RNAP clamp in State I causes $\sigma^3$ detachment from domain 1 (black arrow). The protein linker between $\sigma^3$ and $\sigma^4$ has to accommodate the RNAP clamp opening. The RbpA interaction with both $\sigma^A$ and β' clamp is preserved. **c**, The State II complex superimposed with the PDB entry ID 5TW1 based on the RNAP core domain (β/430-738). In State II, the HelD CO-domain occupies the primary channel and moves even further towards β'-CC, disallowing $\sigma^2$ to bind the β'-clamp. In this situation $\sigma^3$ and $\sigma^4$ hold only on β-domain 1 and β-flap and $\sigma^2$ detaches from the β'-clamp (black arrow). The resulting gap between $\sigma^2$ and β'-clamp would appear filled with the HelD CO-domain.

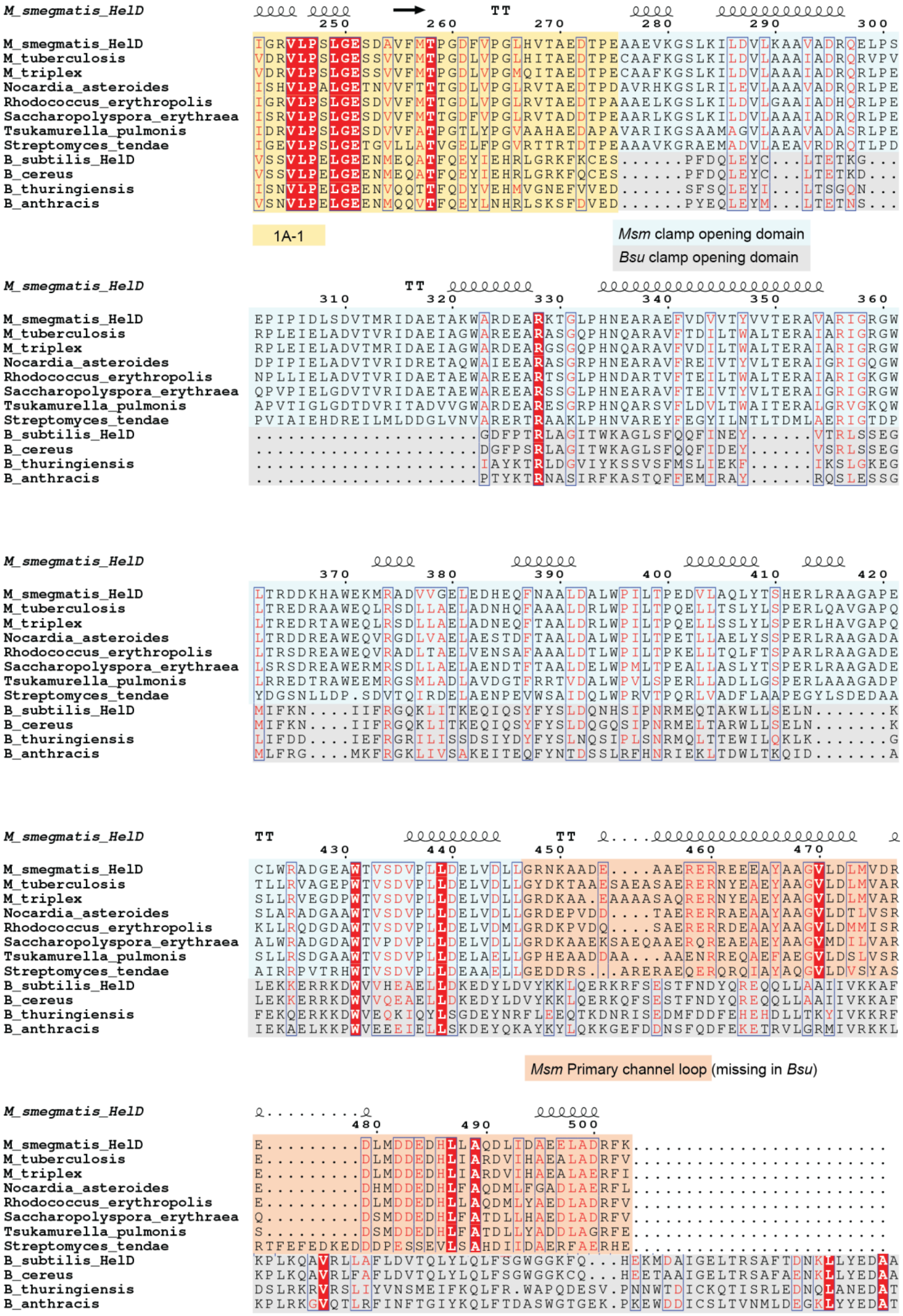
M_smegmatis_HelD
M_tuberculosis
M_triplex
Nocardia_asteroides
Rhodococcus_erythropolis
Saccharopolyspora_erythraea
Tsukamurella_pulmonis
Streptomyces_tendae
B_subtilis_HelD
B_cereus
B_thuringiensis
B_anthracis
1A-1
Msm clamp opening domain
Bsu clamp opening domain
Msm Primary channel loop (missing in Bsu)

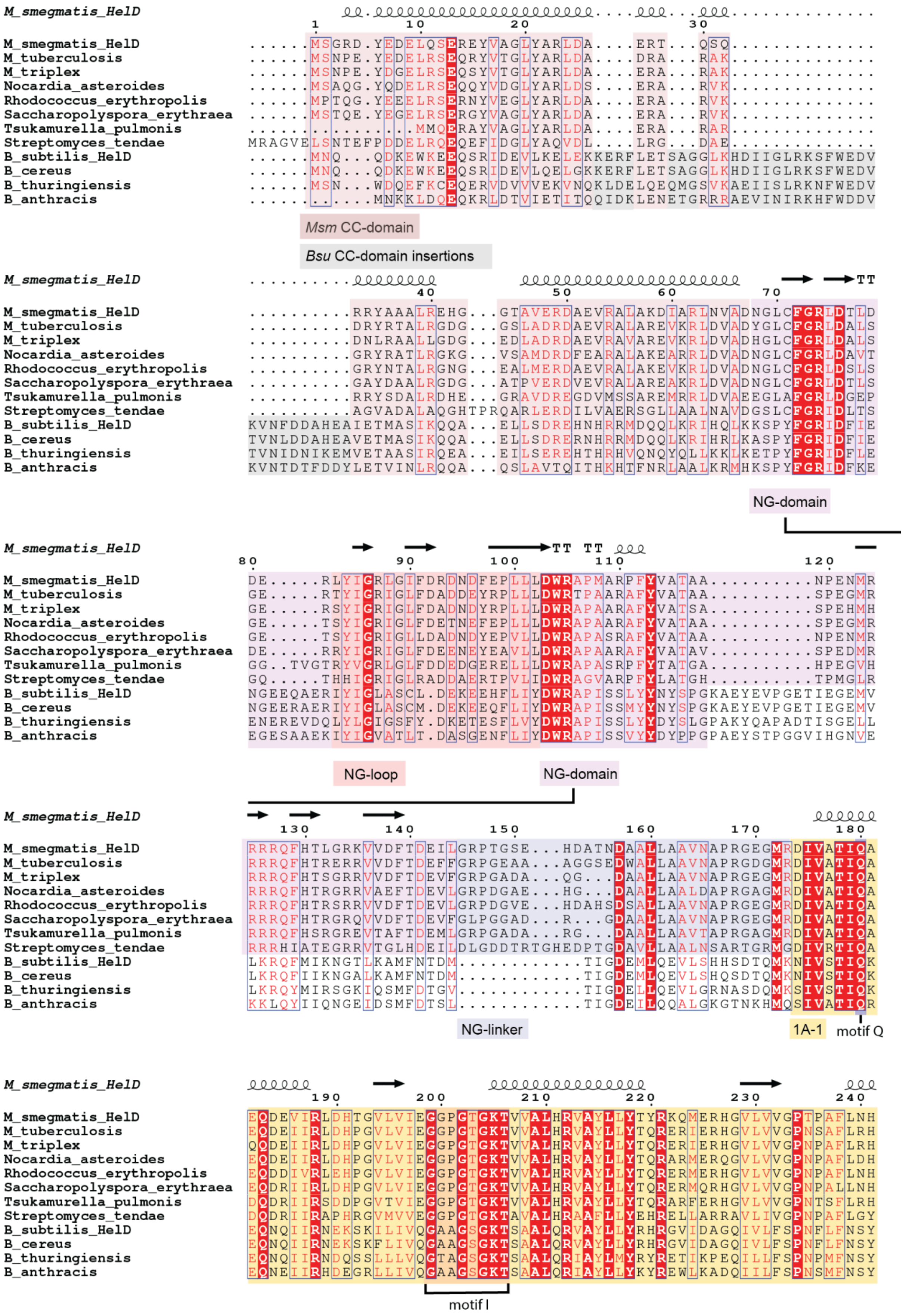
M_smegmatis_HelD
M_tuberculosis
M_triplex
Nocardia_asteroides
Rhodococcus_erythropolis
Saccharopolyspora_erythraea
Tsukamurella_pulmonis
Streptomyces_tendae
B_subtilis_HelD
B_cereus
B_thuringiensis
B_anthracis
Msm CC-domain
Bsu CC-domain insertions
NG-domain
NG-loop
NG-domain
NG-linker
1A-1
motif Q
motif I

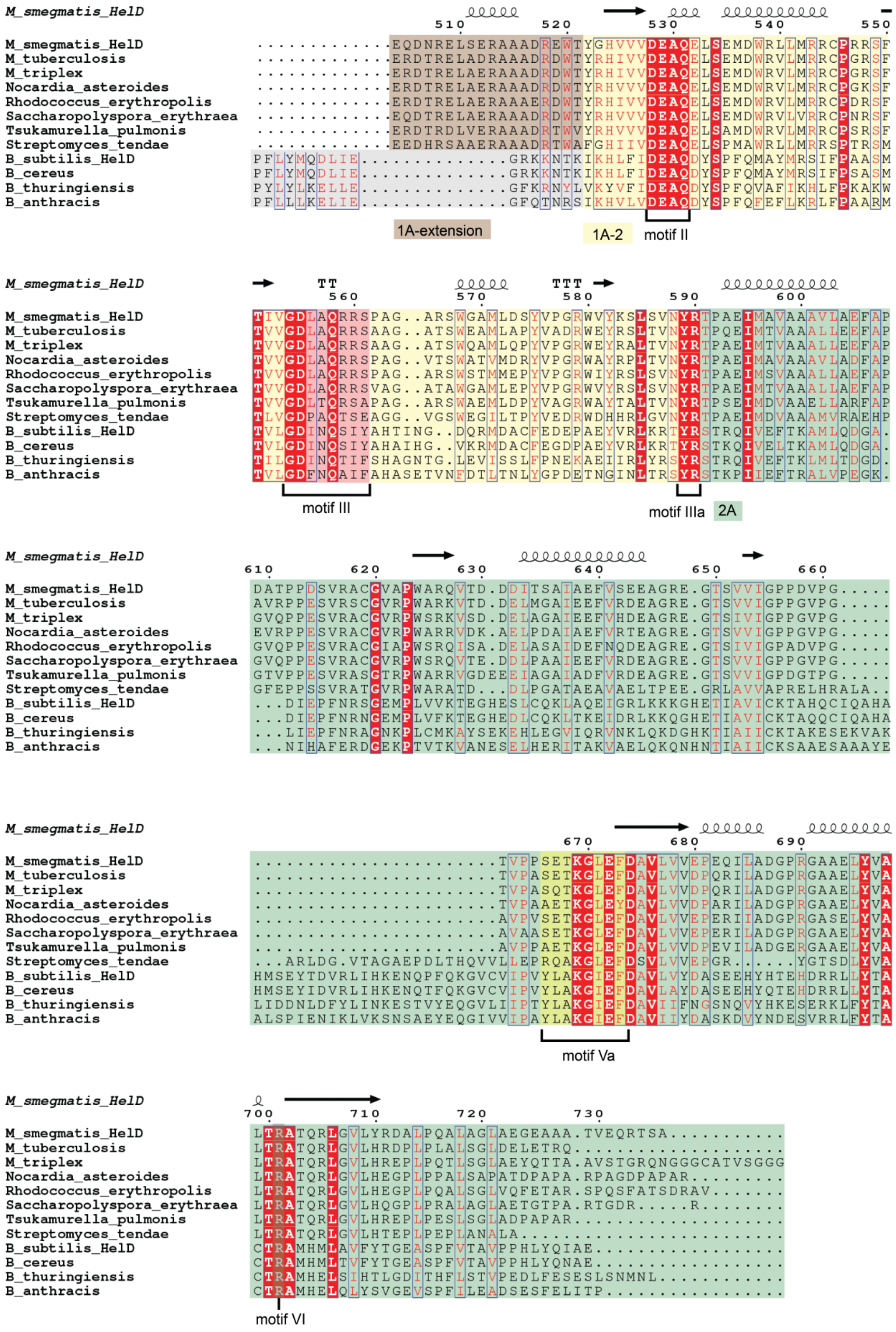
M_smegmatis_HelD
M_tuberculosis
M_triplex
Nocardia_asteroides
Rhodococcus_erythropolis
Saccharopolyspora_erythraea
Tsukamurella_pulmonis
Streptomyces_tendae
B_subtilis_HelD
B_cereus
B_thuringiensis
B_anthracis
1A-extension
1A-2
motif II
motif III
motif IIIa
2A
motif Va
motif VI

**Figure S12: Sequence alignment of HelD homologs**

Curated sequence alignment based on alignment generated by Clustal Omega software[7].GeneBank codes of used sequences: *Msm* WP_003893549.1, *M. tuberculosis*: PLV44927.1; *M. triplex*: CDO88184.1, *Nocardia asteroides*: GAD85771.1, *Rhodococcus erythropolis*: WP_095971734.1, *Saccharopolyspora erythraea*: PFG97077.1, *Tsukamurella pulmonis*: WP_139061895.1; *Streptomyces tendae*: WP_150152972.1, *Bsu* WP_003244180.1, *Bacillus cereus* WP_095971734.1, *B. thuringiensis*: WP_074790911.1, *B. anthracis*: WP_071737252.1.

**Table S1: Hydrogen bonds and salt bridges between HelD RNAP β and β’ subunits**

Interactions up to 4 Å distance according to the PDBe PISA server[6].

**Table S1a: Hydrogen bonds and salt bridges between HelD N-terminal domain (State II) and RNAP β’ subunit**

| # | RNAP β′ subunit | HelD residue |
|---|---|---|
| 1 | D:LYS 775 | H:GLU 27 |
| 2 | D:ASN 809 | H:GLY 43 |
| 3 | D:LYS 820 | H:GLU 48 |
| 4 | D:ARG 865 | H:ASP 50 |
| 5 | D:ARG 757 | H:ASP 96 |
| 6 | D:GLN 778 | H:ARG 34 |
| 7 | D:GLN1008 | H:ARG 49 |
| 8 | D:GLN1146 | H:ARG 49 |
| 9 | D:GLU 751 | H:ARG 93 |
| 10 | D:ASP 779 | H:ARG 93 |
| 11 | D:GLY 762 | H:MET 108 |
| 12 | D:ARG 865 | H:ASP 50 |
| 13 | D:ARG1086 | H:ASP 67 |
| 11 | D:GLU 771 | H:ARG 62 |

**Table S1b: Hydrogen bonds and salt bridges between HelD 1A domain (State II) and RNAP β domain 2 and β’ clamp head**

| # | RNAP β subunit | HelD residue |
|---|---|---|
| 1 | C:LYS 188 | H:THR 521 |
| 2 | C:SER 185 | H:ARG 513 |
| 3 | C:GLU 187 | H:ARG 226 |
| 4 | C:GLU 187 | H:ARG 513 |
| 5 | C:LYS 209 | H:GLU 519 |
| 6 | C:ARG 210 | H:GLU 519 |
| 7 | C:ARG 210 | H:ARG 543 |
| 8 | C:LYS 209 | H:THR 521 |
| 9 | C:ASP 211 | H:ARG 547 |
| | RNAP β′ subunit | |
| 1 | D:VAL1040 | H:GLU 504 |
| 2 | D:LYS1061 | H:GLY 250 |
| 3 | D:ARG1084 | H:GLU 251 |

**Table S1c: Hydrogen bonds and salt bridges between HelD primary channel loop (State I and II) and RNAP β and β' constituents of the primary channel**

| | | |
|---|---|---|
| **State I** | | |
| # | RNAP β' subunit | HelD residue |
| 1 | D:ARG1205 | H:ALA 467 |
| **State II** | | |
| # | RNAP β subunit | HelD residue |
| 1 | C:LYS 184 | H:ASP 500 |
| 2 | C:ARG 456 | H:GLN 490 |
| 3 | C:ARG 464 | H:ASP 491 |
| 4 | C:GLN 605 | H:GLU 484 |
| 5 | C:LYS 875 | H:ASP 483 |
| 6 | C:LYS 883 | H:ASP 483 |
| 7 | C:HIS1026 | H:GLU 484 |
| 8 | C:HIS1026 | H:GLU 484 |
| 9 | C:ARG1058 | H:ASP 479 |
| | RNAP β' subunit | |
| 1 | D:TYR 871 | H:GLU 463 |
| 2 | D:ARG 875 | H:GLU 463 |
| 3 | D:ARG 874 | H:TYR 466 |
| 4 | D:ARG 427 | H:ASP 479 |
| 5 | D:ARG 421 | H:ASP 479 |
| 6 | D:ARG 427 | H:LEU 480 |
| 7 | D:ARG 500 | H:MET 481 |
| 8 | D:GLN 540 | H:MET 481 |
| 9 | D:ALA 542 | H:MET 481 |
| 10 | D:ARG 500 | H:ASP 482 |
| 11 | D:ARG1039 | H:PHE 502 |
| 12 | D:ARG 874 | H:TYR 466 |
| 13 | D:ASP 878 | H:TYR 466 |
| 14 | D:ASP 539 | H:ASP 483 |
| 15 | D:ARG1012 | H:ARG 501 |
| 16 | D:ASP 868 | H:ARG 501 |

**Table S2. Cryo-EM data collection, refinement and validation statistics.**

| | *Msm* HelD-RNAP complex **State I** | *Msm* HelD-RNAP complex **State II** | *Msm* HelD-RNAP complex **State III** |
|---|---|---|---|
| **Deposition** | EMD-10996, PDB ID 6YXU | EMD-11004, PDB ID 6YYS | EMD-11026, PDB ID 6Z11 |
| **Data collection and processing** | | | |
| Magnification | 165,000 | 165,000 | 165,000 |
| Voltage (kV) | 300 | 300 | 300 |
| Electron exposure (e–/Å$^2$) | 40-50 | 40-50 | 40-50 |
| Defocus range (μm) | 0.7-3.3 | 0.7-3.3 | 0.7-3.3 |
| Pixel size (Å) | 0.8311 | 0.8311 | 0.8311 |
| Symmetry imposed | C1 | C1 | C1 |
| Initial particle images (no.) | 1,560,500 | 1,560,500 | 1,560,500 |
| Final particle images (no.) | 185,400 | 173,500 | 119,100 |
| Map resolution (Å) | 3.08 | 3.08 | 3.47 |
| FSC threshold | 0.143 | 0.143 | 0.143 |
| Map resolution range (Å) | 3.08-5.90 | 3.02-5.90 | 3.29-5.90 |
| **Refinement** | | | |
| Initial model used (PDB code) | 6F6W[8] | 6F6W | 6F6W |
| Model resolution (Å) | 3.2 | 3.2 | 3.5 |
| FSC threshold | 0.5 | 0.5 | 0.5 |
| Model resolution range (Å) | 3.09-5.90 | 3.02-5.90 | 3.05-5.90 |
| Map sharpening *B* factor (Å$^2$) | -78.53 | -81.37 | -85.45 |
| Model vs map cross correlation | 0.81 | 0.79 | 0.81 |
| Model composition | | | |
| Non-hydrogen atoms | 27791 | 27930 | 23948 |
| Protein residues | 3583 | 3597 | 3077 |
| Nucleotide residues | 0 | 0 | 0 |
| Ligands | 3 | 3 | 3 |
| *B* factors (Å$^2$) | | | |
| Protein | 40.27 | 32.39 | 34.47 |
| Ligand | 61.69 | 47.49 | 46.56 |
| R.m.s. deviations from ideal | | | |
| Bond lengths (Å) | 0.006 | 0.005 | 0.005 |
| Bond angles (°) | 0.672 | 0.656 | 0.610 |
| Validation | | | |
| MolProbity score | 2.03 | 2.00 | 2.01 |
| Clashscore | 9.28 | 9.18 | 7.94 |
| Poor rotamers (%) | 0.00 | 0.00 | 0.04 |
| Ramachandran plot | | | |
| Favored (%) | 90.54 | 91.15 | 89.03 |
| Allowed (%) | 9.43 | 8.82 | 10.97 |
| Disallowed (%) | 0.03 | 0.03 | 0 |

**Table S3: Data collection and refinement statistic of the *B. subtilis* HelD C-terminal domain. Values in parentheses refer to the highest resolution shell.**

| PDB code | 6VSX |
|---|---|
| **Data collection** | |
| X-ray source | Rigaku MicroMax 007 HF |
| Wavelength (Å) | 1.54178 |
| No. if oscillation images | 1080 |
| Total oscillation angle | 1080 |
| Delta phi (°) | 1 |
| Crystal to detector distance (mm) | 50 |
| Average mosaicity (°) | 1.4 |
| Space group | $C2_1$ |
| Cell dimensions | |
| $a$ (Å) | 106.96 |
| $b$ (Å) | 38.81 |
| $c$ (Å) | 44.43 |
| β (º) | 101.45 |
| Resolution (Å) | 25.0 – 2.0 |
| No. of all observed reflections | 245,968 |
| No. of unique reflections | 11,905 |
| Average redundancy | 20.7 (14.1) |
| Completeness (%) | 96.7 (72.0) |
| $I/\sigma(I)$ | 60.1 (14.3) |
| Wilson B-factor (Å$^2$) | 21.87 |
| R-merge | 0.044 (0.206) |
| CC1/2 | (0.991) |
| CC* | (0.998) |
| **SAD Phasing (S and P)** | |
| Number of sites | 10 (S) and 1 (P) |
| Figure of Merit | 0.296 |
| **Refinement** | |
| Resolution (Å) | 25.0 – 2.0 |
| No. of reflections used in refinement | 11,869 (1,186) |
| $R_{work}$ | 0.1723 (0.1756) |
| $R_{free}$ | 0.2014 (0.2393) |
| No. of atoms | 1,382 |
| macromolecules | 1,268 |
| ligands | 5 |
| solvent | 109 |
| No. of protein residues | 159 |
| RMS deviations from ideal | |
| bond lengths (Å) | 0.007 |
| bond angles (°) | 0.80 |
| Clashscore (Molprobity) | 5.92 |
| Ramachandran plot, residues | |
| in favored region (%) | 98.06 |
| outliers (%) | 0.0 |
| Average B-factor (Å$^2$) | 25.2 |
| Macromolecules (Å$^2$) | 24.6 |
| Ligands (Å$^2$) | 30.5 |
| Solvent (Å$^2$) | 32.9 |

**Supplementary references**


1. Zheng, S. Q. *et al.* MotionCor2: anisotropic correction of beam-induced motion for improved cryo-electron microscopy. *Nat Methods* **14**, 331-332, doi:10.1038/nmeth.4193 (2017).
2. Rosenthal, P. B. & Henderson, R. Optimal determination of particle orientation, absolute hand, and contrast loss in single-particle electron cryomicroscopy. *J Mol Biol* **333**, 721-745 (2003).
3. Zivanov, J. *et al.* New tools for automated high-resolution cryo-EM structure determination in RELION-3. *Elife* **7**, doi:10.7554/eLife.42166 (2018).
4. Zivanov, J., Nakane, T. & Scheres, S. H. W. Estimation of high-order aberrations and anisotropic magnification from cryo-EM data sets in RELION-3.1. *IUCrJ* **7**, 253-267, doi:10.1107/S2052252520000081 (2020).
5. Lee, J. Y. & Yang, W. UvrD helicase unwinds DNA one base pair at a time by a two-part power stroke. *Cell* **127**, 1349-1360, doi:10.1016/j.cell.2006.10.049 (2006).
6. Krissinel, E. & Henrick, K. Inference of macromolecular assemblies from crystalline state. *J Mol Biol* **372**, 774-797, doi:10.1016/j.jmb.2007.05.022 (2007).
7. Madeira, F. *et al.* The EMBL-EBI search and sequence analysis tools APIs in 2019. *Nucleic Acids Res* **47**, W636-W641, doi:10.1093/nar/gkz268 (2019).
8. Kouba, T. *et al.* The Core and Holoenzyme Forms of RNA Polymerase from Mycobacterium smegmatis. *J Bacteriol* **201**, doi:10.1128/JB.00583-18 (2019).
